# Amphiphilic Peptide Fusion Promotes Endocytic Uptake of Nanodiscs

**DOI:** 10.64898/2026.05.08.723726

**Authors:** Brandon S. Pizarro, Trevin G. Reinhardt, Jason A. Semenske, Zekai Ji, Ciara O. Jacobs, Wade Zeno

**Affiliations:** Mork Family Department of Chemical Engineering and Materials Science, University of Southern California, Los Angeles, California 90089, United States; Department of Physics, Río Hondo College, Whittier, California, 90601, United States

## Abstract

A major limitation across nanoparticle delivery platforms is sequestration within endosomal compartments, which restricts access to intracellular targets despite efficient cellular uptake. Here, we show that peptide architecture can be used to control intracellular trafficking and reduce endosomal accumulation in lipid-protein nanocarriers. Specifically, we fuse R6W3 (RRWWRRWRR), an amphipathic cell penetrating peptide, to the N- or C- terminus of the nanodisc scaffold proteins and systematically evaluate its impact on membrane interactions and cellular behavior. Structural and biophysical characterization confirms that R6W3 incorporation preserves nanodisc assembly and protein-lipid interactions, enabling direct attribution of functional differences to peptide-driven interfacial effects. R6W3-functionalized nanodiscs exhibit enhanced binding and cellular uptake, with N-terminal fusion producing the strongest interfacial interactions. In live cells, R6W3-functionalization increases endocytic activity, evidenced by increased formation of clathrin-coated pits and intracellular colocalization with clathrin-coated vesicles. Notably, R6W3-funtionalized nanodiscs display reduced accumulation in early endosomes relative to unmodified nanodiscs, indicating decreased endosomal sequestration following endosomal uptake. These trafficking differences translate to functional outcomes, as doxorubicin-loaded, R6W3-functionalized nanodiscs achieve greater cytotoxicity than unmodified controls at equivalent concentrations. Together, these results establish peptide architecture as a design parameter for controlling intracellular trafficking and overcoming endosomal bottlenecks, providing a broadly applicable strategy for improving nanocarrier- based delivery systems.

## INTRODUCTION

Delivery platforms that mimic endogenous lipid carriers are of growing interest for cellular delivery applications due to their inherent biocompatibility^1, 2^. In particular, high-density lipoprotein (HDL) platforms have attracted attention for their ability to engage native cellular pathways and facilitate receptor-mediated uptake^3, 4^. Small, therapeutic molecules, such as doxorubicin (DOX), can be incorporated into HDL particles and delivered to cells^5^. However, native HDL particles are compositionally heterogeneous and undergo continuous lipid exchange and remodeling in biological environments^6^, limiting precise control over surface functionalization and cargo presentation.

To address these limitations, HDL mimetic nanoparticles, commonly referred to as nanodiscs, have emerged as a structurally defined and biologically inspired platform^7, 8^. Nanodiscs consist of a planar phospholipid bilayer encircled by membrane scaffold proteins (MSPs), amphipathic derivatives of apolipoprotein A-I (ApoAI) that self-assemble into a helical belt around the lipid bilayer^9, 10^. This architecture enables precise control over particle size and lipid composition while maintaining compatibility with biological membranes. Importantly, MSP scaffolds tolerate extensive genetic modification, enabling incorporation of functional motifs without disrupting nanodisc structure^11, 12^.

One widely used strategy to enhance nanodisc uptake is the incorporation of cell-penetrating peptides (CPPs) through genetic fusion to the MSP ^13–16^ . Classical cationic CPPs, including arginine-rich sequences such as TAT, penetratin, and polyarginine, promote membrane association primarily through electrostatic interactions^17–19^. However, these peptides exhibit heterogenous internalization behavior^20, 21^, with reported contributions from multiple pathways including macropinocytosis, clathrin mediated endocytosis (CME) and direct translocation^22–26^. As a result, the extent to which peptide architecture govern nanodisc uptake and routing through specific cellular pathways remains unclear.

R6W3 (RRWWRRWRR) is a synthetic amphipathic CPP that integrates arginine-mediated electrostatic interactions with tryptophan-driven hydrophobic engagement^27, 28^. Unlike many classical cationic CPPs, R6W3 adopts an amphipathic l1-helical conformation upon interacting with lipid membranes^29^ and has been shown to induce lipid remodeling in model membranes^30^. These properties suggest a distinct mode of interaction with cellular membranes. Notably, the asymmetric distribution of arginine and tryptophan residues along the peptide sequence introduces directional amphiphilicity, raising the possibility that N- versus C- terminal fusion to MSPs may differentially influence nanodisc cell interactions. Despite the extensive characterization of R6W3 behavior^31–33^, to the best of our knowledge, R6W3 has yet to be incorporated into nanoparticle-based delivery systems.

Here, we incorporate R6W3 into MSP-based nanodiscs and investigate how this amphiphilic peptide fusion modulates nanodisc interactions with lipid bilayers and cellular uptake. Using quantitative fluorescence measurements and live cell imaging, we demonstrate that R6W3 enhances binding to lipid vesicles and cellular uptake, with N-terminal fusion producing stronger synthetic membrane association than C- terminal fusion while both terminal constructs comparably increase cellular internalization relative to unmodified nanodiscs. R6W3-functionalized nanodiscs elicit increased endocytic activity and colocalize with clathrin-coated vesicles, directly establishing CME as the dominant internalization pathway. Beyond CME-mediated entry, R6W3-funcitonalized nanodiscs exhibit reduced accumulation in early endosomes, suggesting decreased sequestration within these compartments and increased access to intracellular space. This shift in intracellular trafficking correlates with enhanced cytotoxic potency of DOX-loaded nanodiscs, indicating more effective intracellular delivery of functional cargo. Together, these findings establish peptide architecture as a design parameter for controlling nanodiscs endocytic routing and improving intracellular cargo delivery.

## RESULTS

### Synthesis and characterization of R6W3-functionalized nanodiscs

To evaluate R6W3 functionalization of nanodiscs (NDs), we engineered two peptide-MSP fusion constructs with R6W3 fused to either the C-terminus (C-term) or N-terminus (N-term) and compared their membrane and cellular interactions to unmodified MSP as a control (Fig. 1A, Table S1). Here, we used MSP1D1, a commonly employed nanodisc scaffold protein derived from the amphipathic helices of ApoAI^9^, with hexahistidine tags included for purification. Successful incorporation of R6W3 into the MSP constructs was confirmed with SDS PAGE (Fig. 1B), where the C-term and N-term variants exhibited comparable molecular weights that were slightly larger than that of the unmodified MSP. Using these protein constructs, NDs were assembled via a detergent based self-assembly protocol (Fig. 1C). Dynamic light scattering (DLS) was used to monitor the self-assembly process (Fig. 1D), with all ND formulations converging to populations with an average hydrodynamic diameter of approximately 20 nm after purification. The precise size of NDs was determined via transmission electron microscopy (TEM), where ND diameters were confirmed to be between 18-20 nm on average (Fig. 1E). The consistent size between unmodified and R6W3-functionalized NDs indicates that the attachment of R6W3 does not inhibit the ND self-assembly procedure nor significantly alter particle dimensions.

**Figure 1:**
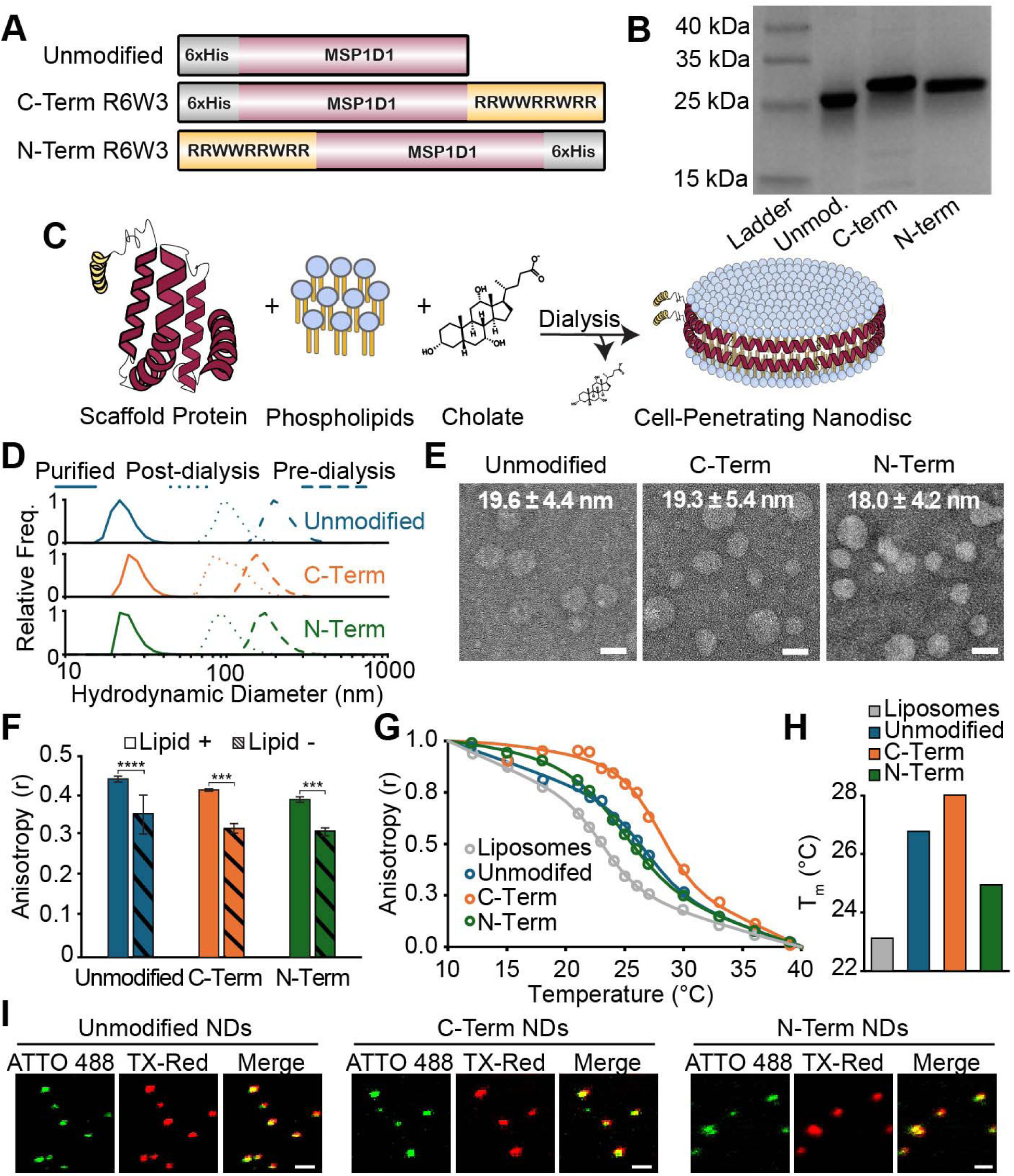
Assembly and characterization of R6W3-functionalized NDs. (A) Schematic of MSP1D1-derived constructs. (B) SDS-PAGE images of purified, MSP1D1-derived proteins constructs. (C) Schematic of ND self-assembly procedure. (D) DLS-measured hydrodynamic diameter distributions during ND synthesis and after purification. (E) Representative TEM images and corresponding size distribution of NDs. (F) Fluorescent anisotropy of scaffold protein in the presence (+) or absence (-) of lipids. (G) Temperature-dependent fluorescence anisotropy profiles of DPH within lipid bilayers. (H) Phase transition temperatures (T_m_) of lipid bilayers in either liposomes or NDs, as determined by DPH anisotropy measurements. (I) Representative fluorescence images showing colocalization of lipids (Texas-Red) and proteins (ATTO 488) within NDs. Data are presented as mean ± SEM. Statistical significance was determined by two tailed unpaired Student’s t-test (* = p < .05, ** = p < .01, *** = p < .001, **** = p < .0001). Scale bars: (E) 20 nm; (I) 2 µm.

Next, a series of experimental measurements were performed to confirm intact protein-lipid interactions within the NDs (Figs. 1F-I). Using the intrinsic fluorescence of tryptophan residues within MSPs, we performed anisotropy measurements on lipid-containing and lipid-free samples (Fig. 1F). In these specific measurements, larger fluorescence anisotropy values correspond to slower rotational speeds of the fluorophore and therefore, a larger effective size of the fluorophore-containing moiety^34^. The higher anisotropy values in lipid-containing samples across all formulations indicate that MSPs are attached to lipids in ND complexes, rather than existing as individual proteins in solution.

Fluorescence anisotropy measurements were also performed using the lipid bilayer-partitioning fluorescent probe 1,6-Diphenyl-1,3,5-hexatriene (DPH) (Fig. 1G). Here, fluorescence anisotropy measurements represent the rotational freedom of DPH within DMPC bilayers. As the samples are heated beyond the melting temperature of DMPC (i.e., 23°C in liposomal form^35^), lipid packing is disrupted and DPH rotational speed increases, resulting in a decrease in anisotropy (Fig. 1G). The elevated phase transition temperatures of DMPC in ND formulations (Figs. 1H) further demonstrate protein-lipid association, as MSPs are known to order lipids within NDs^7, 36^, increasing the energetic threshold for melting.

Lastly, colocalization between lipids and proteins within NDs was observed using confocal fluorescence microscopy (Fig. 1I). By fluorescently labeling MSPs with ATTO 488 NHS-ester and including Texas Red DHPE in DMPC bilayers, individual NDs were imaged and clear overlap of fluorescence channels was observed for each construct. Together, these results indicate the feasibility of synthesizing stable NDs with R6W3-MSP fusion constructs.

### R6W3-functionalization of nanodiscs facilitates membrane binding

We next used giant unilamellar vesicles (GUVs) to determine whether R6W3-functionalized NDs associate with lipid bilayers (Fig. 2). Confocal fluorescence microscopy was used to image GUVs at their equatorial plane, either in the presence or absence of NDs (Fig. 2A). GUVs were fluorescently labeled via incorporation of ATTO 647N DPPE, while NDs were labeled via Texas Red DHPE (lipids) and ATTO 488 NHS-ester (scaffold proteins). Upon incubation with GUVs, R6W3-functionalized NDs exhibited increased membrane-associated fluorescence relative to unmodified nanodiscs, as determined by radial scan profiles (Fig. 2B). Quantitative analysis demonstrated a concentration-dependent increase in membrane-associated fluorescence for R6W3-functionalized nanodiscs. In particular, the N-term variant demonstrated significantly enhanced membrane association across both the lipid and protein fluorescence channels (Figs. 2C-D). Similar binding behavior was observed using enhanced green fluorescent protein (eGFP) as a control, where R6W3 was either C- or N-terminally fused to the protein (Figs. 2E, S1, and Table S1). Together, these results indicate that R6W3-functionalization imparts membrane binding abilities to NDs in a terminal-dependent manner.

**Figure 2:**
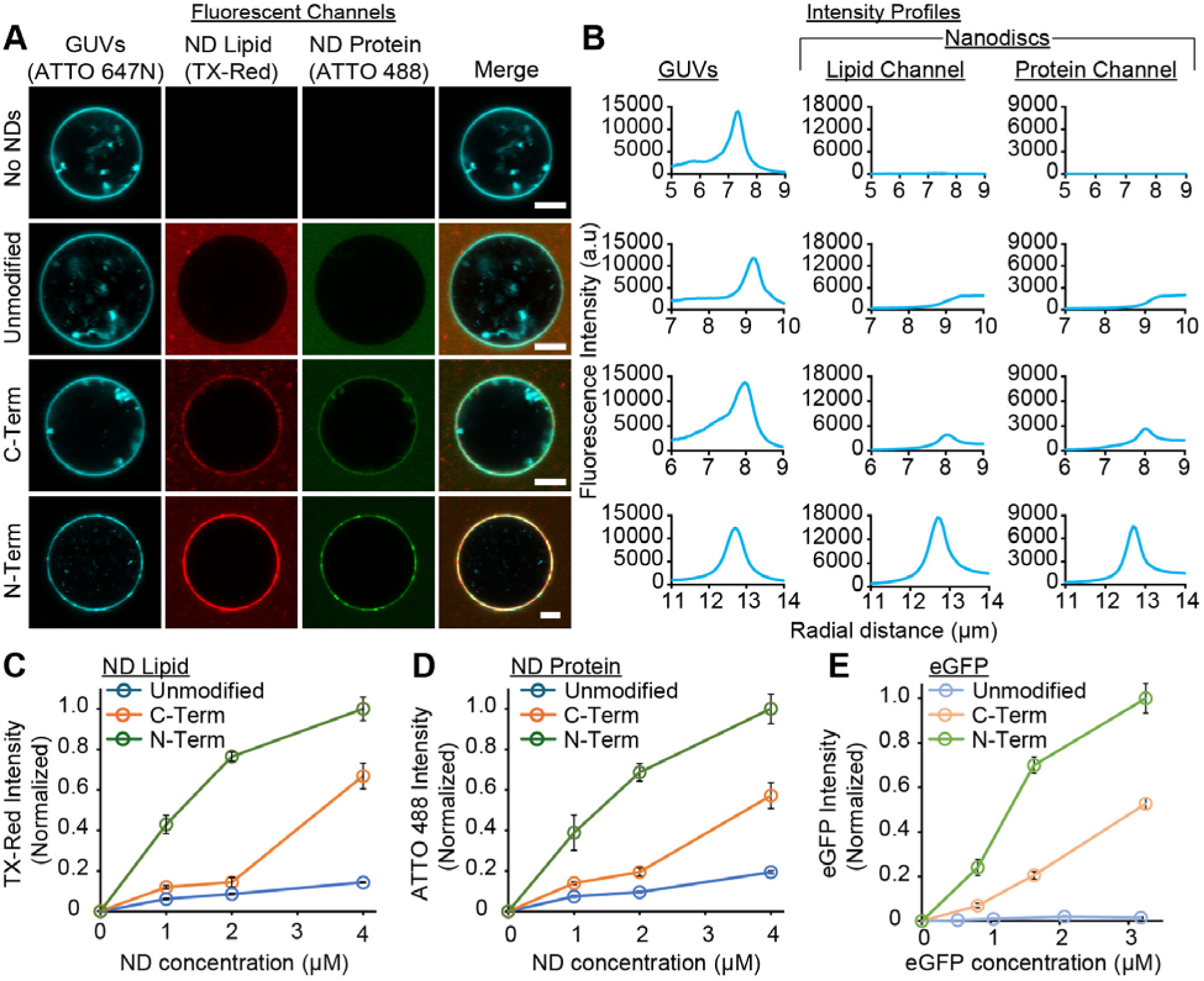
R6W3 functionalization enhances membrane association with model membranes. (A) Confocal fluorescence images of GUVs (49% DOPS, 49% DOPC, 1% ATTO 647N DPPE, 1% DPPE-Cap-biotin) incubated with NDs containing Texas-Red labeled lipids and ATTO 488 labeled MSP1D1. Merged images show NDs association with GUV membranes. (B) Radial fluorescence intensity profiles of GUV and corresponding ND lipid and ND protein. (C-D) Quantification of membrane associated signal from radial profiles for NDs in (C) the lipid fluorescence channel and (D) the protein fluorescence channel as a function of ND concentration. (E) Membrane association of eGFP constructs modified analogous to NDs. Data are presented as mean ± SEM. Scale bar: 5 µm.

### Impact of R6W3-functionalization on nanodisc association to live cells

After confirming the ability of R6W3-functionalized NDs to bind lipid bilayers, we next quantified their ability to associate with cells (Fig. 3). To determine the impact of ND modification, we first utilized epifluorescence microscopy to visualize association with SUM159 cells (Figs. 3A-B). Cells incubated with free ATTO 488 dye or unmodified, fluorescent nanodiscs showed sparse fluorescent overlap with cellular regions, whereas incubation with fluorescent N-term nanodiscs resulted in discernible fluorescence within cellular regions in the form of punctate structures (Fig. 3A).

**Figure 3:**
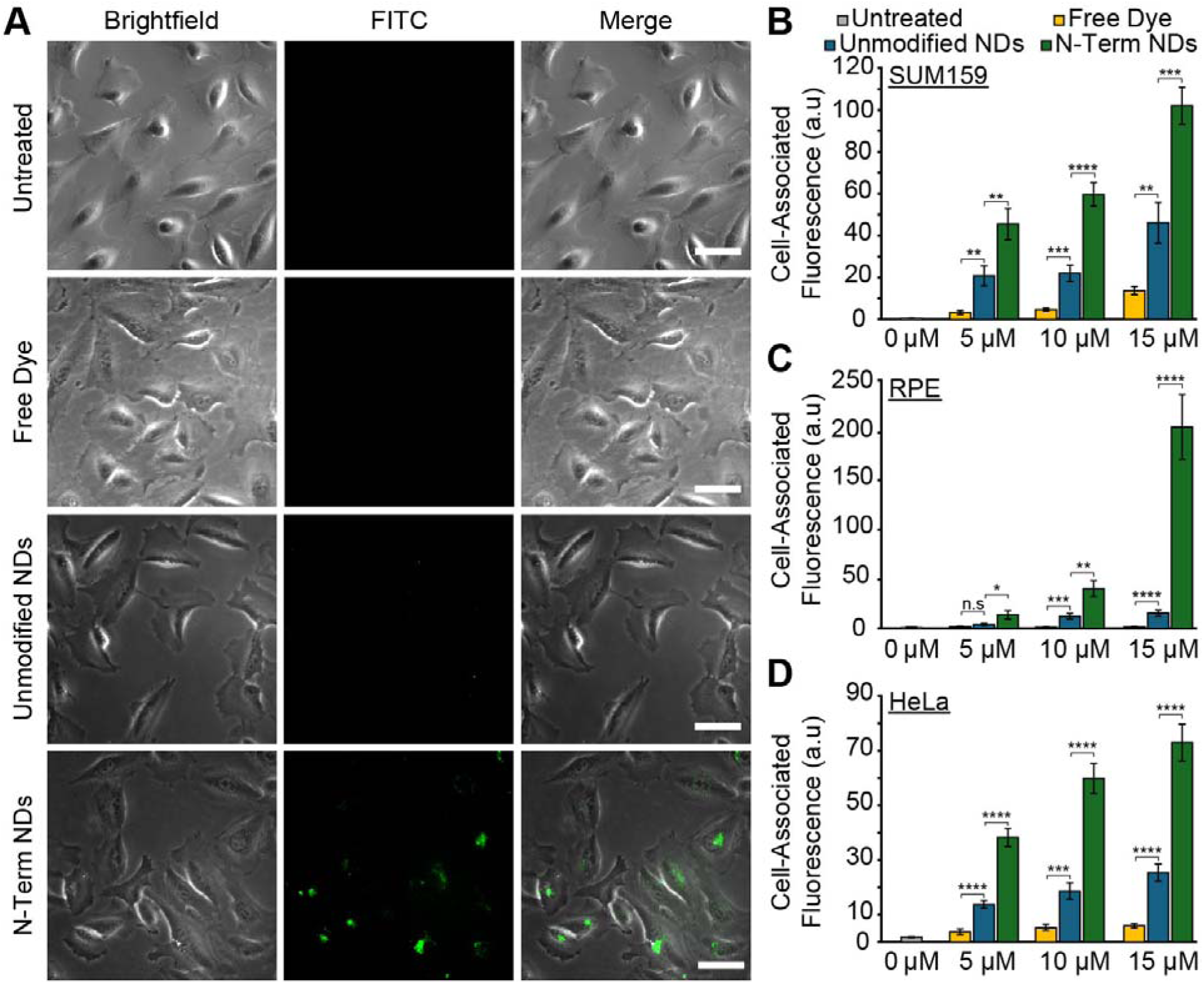
R6W3-functionalized NDs increase cellular association across multiple cell types. (A) Representative brightfield and fluorescence images of SUM159 cells following incubation with fluorescent NDs, free ATTO 488 dye, or untreated controls. (B-D) Quantification of cell-associated fluorescence in (B) SUM159, (C) RPE, and (D) HeLa cells as a function of ND concentration. Data are presented as mean ± SEM. Statistical significance was determined by two tailed unpaired Student’s t-test (* = p < .05, ** = p < .01, *** = p < .001, **** = p < .0001, n.s = not significant). Scale bars: 50 µm.

The cell-associated fluorescence signal, calculated from equation 3 (see methods), demonstrated a monotonic response within each condition, with N-term NDs eliciting the highest signal (Fig. 3B). Similar results were observed for fluorescent C-term NDs (Fig. S2), with comparable cell-associated fluorescence to N-term NDs. Qualitatively similar cell-association trends were also observed in retinal pigment epithelial (RPE) cells (Figs. 3C, S3) and HeLa cells (Figs. 3D,S4).

Control experiments with N- and C-terminally fused R6W3 in eGFP were performed for all cell types listed above (Figs. S5-S7). While R6W3 fusion enhanced cellular association for both eGFP constructs relative to unmodified eGFP, N-terminal fusion elicited stronger cellular association than the C-terminal variant. Additionally, modified eGFP constructs exhibited diffuse fluorescence throughout cellular regions, as opposed to the punctate structures observed with modified NDs.

### Impact of R6W3-functionalization on cellular internalization of nanodiscs

Following the observed increase in cellular association, we next examined whether R6W3-functionalized NDs were internalized by cells, as epifluorescence microscopy cannot distinguish between plasma membrane adsorption and intracellular localization. To perform this analysis, we used confocal fluorescence microscopy, acquiring optical sections through cells (Fig. 4A).

**Figure 4:**
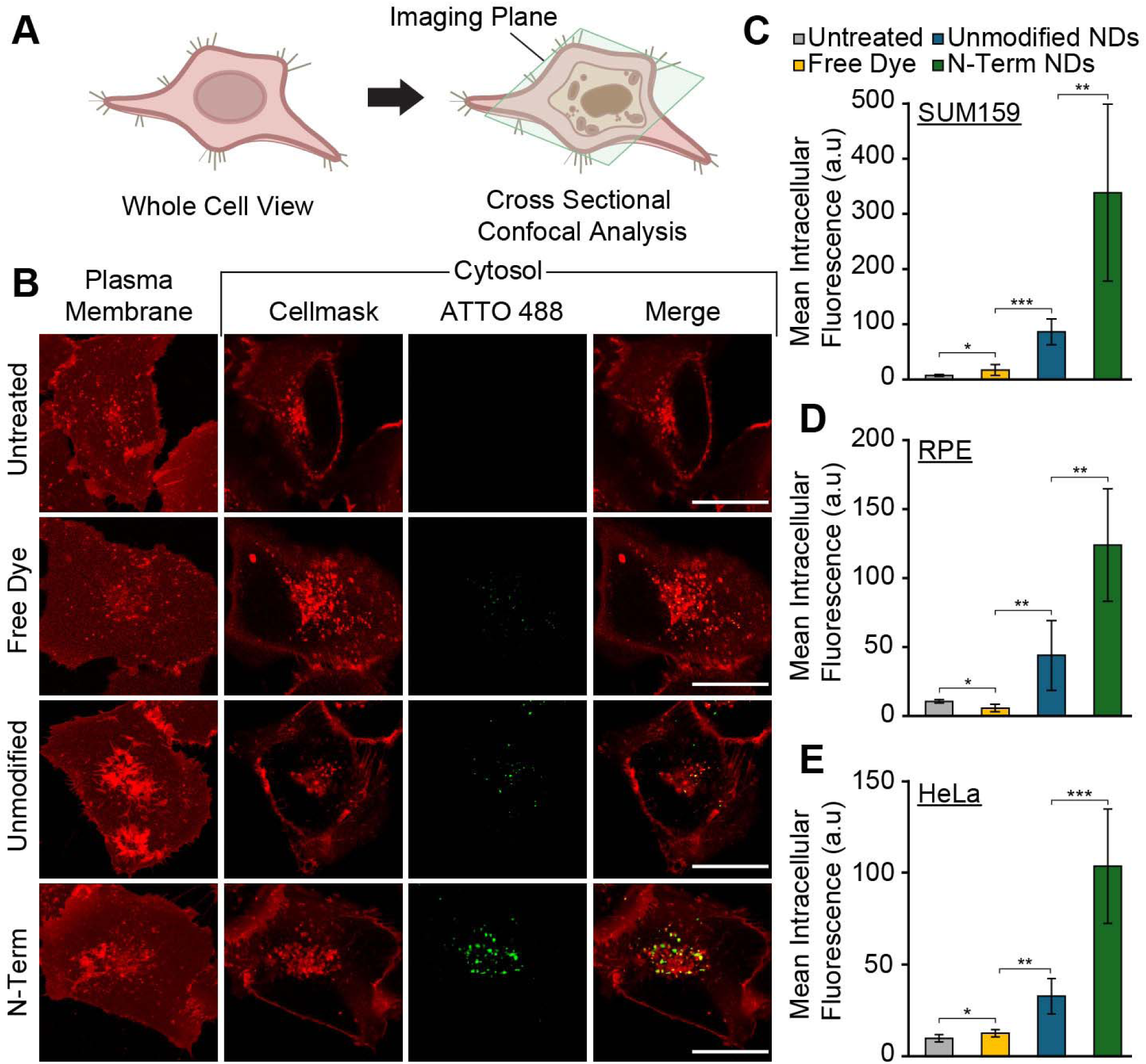
R6W3-funtcionalized NDs undergo cellular internalization and accumulate intracellularly. (A) Schematic illustrating optical sectioning by confocal microscopy used to distinguish membrane-associated and intracellular ND signal. (B) Representative confocal images of SUM159 cells stained with CellMask™ (membrane/cytosol, red) and incubated with fluorescent NDs (green). (C-E) Quantification of intracellular fluorescence in (C) SUM159, (D) RPE, and (E) HeLa cells at 10 µM ND concentration. Data are presented as mean ± SD. Statistical significance was determined by two tailed unpaired Student’s t-test (* = p < .05, ** = p < .01, *** = p < .001, **** = p < .0001). Scale bar: 25 µm.

SUM159 cells were stained with CellMask™ dye and imaged at both the plasma membrane and within the cytosolic region following incubation with free ATTO 488 or fluorescently labeled NDs (Fig 4B). The plasma membrane was used as a reference plane to distinguish surface-associated signal from intracellular localization. Conditions with free dye or unmodified NDs yielded sparse fluorescent signal within cells, whereas N-term NDs exhibited visibly stronger fluorescence signal with punctate structures. The mean intracellular fluorescence intensity was highest for SUM159 cells incubated with N-term NDs (Fig. 4C). Qualitatively similar results were observed for RPE cells (Figs. 4D, S9) and HeLa cells (Figs. 4E, S10). C-term NDs exhibited greater cellular uptake than unmodified NDs, but comparable or lower uptake than N-term NDs across all cell lines (Figs S8-S10).

### Trafficking of R6W3-functionalized nanodiscs in live cells

To gain insight into the internalization mechanism of R6W3-functionalized NDs, we examined their cellular interactions in SUM159 cells expressing fluorescently tagged AP2, which allowed monitoring of endocytic pits and vesicles during CME (Figs. 5A-D). At the plasma membrane surface, incubation with fluorescent N-term NDs yielded the highest fluorescence intensity (Fig 5A), suggesting that N-term NDs exhibit the highest degree of binding to the plasma membrane. Consistent with this elevated binding, N-term NDs also significantly increased the density of clathrin-coated pits on the plasma membrane (Figs. 5B-C). C-term NDs also increased plasma membrane binding relative to unmodified NDs and increased the density of endocytic pits (Fig. S11). Intracellular imaging confirmed that cells exposed to N-term NDs contained an elevated density of clathrin-coated vesicles within the cytosol (Fig. 5D). N-term ND fluorescence was distinctly colocalized with these cytosolic clathrin-coated vesicles. This increased CME activity and colocalization with clathrin-coated vesicles were also observed in cells incubated with C-term NDs (Fig. S11).

**Figure 5:**
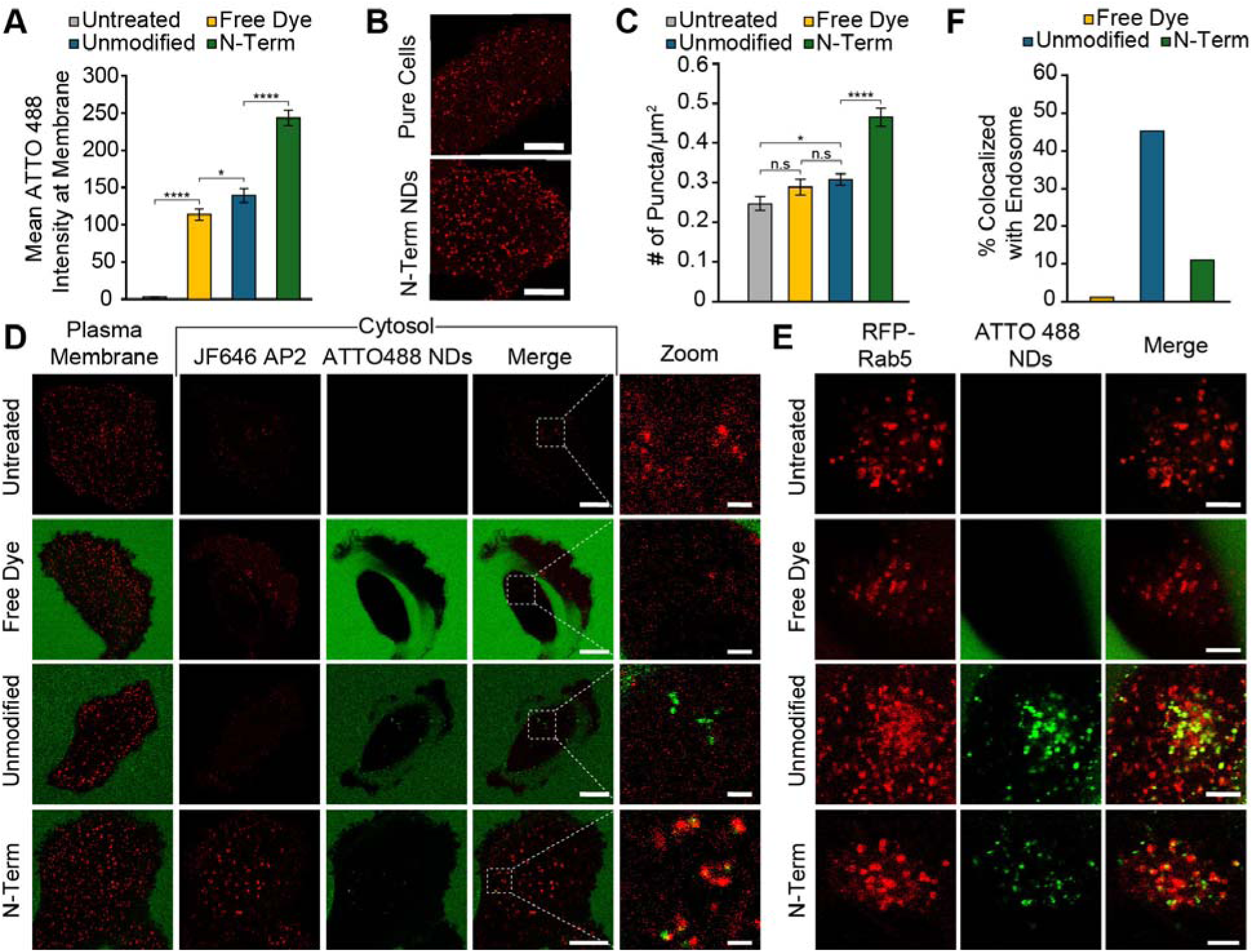
R6W3-functionalized NDs increase endocytic activity and exhibit decreased accumulation in early endosomes within SUM159 cells. (A) Quantification of plasma membrane-associated ATTO 488 fluorescence in SUM159 cells. (B) Representative fluorescence images highlighting increased density of punctate structures (i.e., clathrin-coated pits) on the plasma membrane following treatment with N-Term NDs. (C) Quantification of clathrin-coated pit density. (D) Representative confocal images of SUM159 expressing AP2-HaloTag labeled with JF646 (red) and incubated with fluorescent NDs (green). Insets show magnified regions of interest. (E) Representative confocal images of intracellular ATTO 488 signal and early endosomes labeled with Rab5-RFP. (F) Quantification of ATTO 488 colocalization with endosomes across treatment conditions. Data are presented as mean ± SEM. Statistical significance was determined by two tailed unpaired Student’s t-test (* = p < .05, ** = p < .01, *** = p < .001, **** = p < .0001). Scale bars: (B) 10 µm; (D) 10 µm (main images) and 1 µm (zoom); (E) 5 µm.

We next examined ND association with early endosomes, the canonical destination following CME. Early endosomes were visualized using fluorescent Rab5 (Fig. 5E). Substantial intracellular fluorescence was observed for both types of NDs. However, by visual inspection, the unmodified variant exhibited a higher degree of colocalization with early endosomes than the N-term formulation. Quantitative analysis confirmed this observation (Fig. 5F). C-term NDs also displayed reduced colocalization with early endosomes (Fig. S12). Together, these results demonstrate that R6W3-functionalized NDs enhance CME activity in cells while exhibiting decreased accumulation in early endosomes.

### Drug delivery with R6W3-functionalized nanodiscs

To assess the efficacy of R6W3-functionalized NDs in drug delivery, we modified our ND self-assembly protocol to incorporate doxorubicin (DOX) (Fig. 6A), a chemotherapeutic that induces cell death by intercalating DNA. TEM confirmed the formation of disc-shaped nanoparticles with mean diameters ranging from approximately 11 to 13 nm (Fig. 6B, Fig. S13 for C-term NDs). Absorbance spectroscopy confirmed that each ND contained approximately two DOX molecules on average across all formulations (see methods).

**Figure 6:**
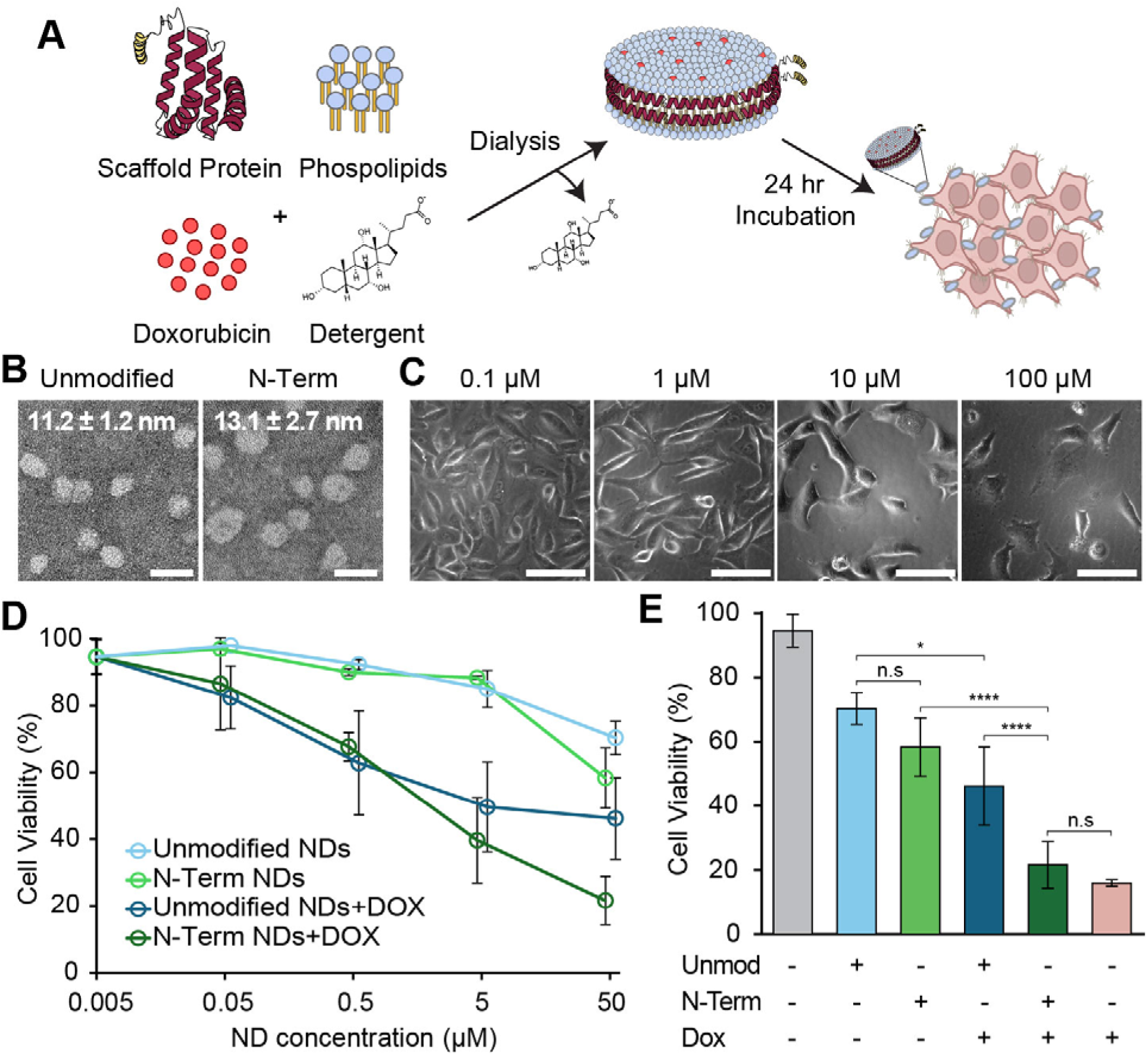
DOX-loaded, R6W3-functionalized NDs exhibit enhanced cytotoxicity in SUM159 cells. (A) Schematic of ND self-assembly with DOX incorporation. (B) Representative TEM images of DOX-containing NDs. TEM size data are presented as mean ± SEM. (C) Brightfield micrographs of SUM159 cells following treatment with increasing concentrations of DOX-loaded N-term NDs. (D) Cell viability as a function of ND concentration. (E) Comparison of cell viability across treatment conditions at the highest DOX concentration tested. Cell viability data are presented as mean ± SD. Statistical significance was determined by two tailed unpaired Student’s t-test (* = p < .05, ** = p < .01, *** = p < .001, **** = p < .0001). Scale bars: (B) 20 nm; (C) 100 µm.

To quantify DOX delivery, SUM159 cells were incubated with either DOX-free or DOX-containing NDs and cell viability was calculated (Figs. 6C-D). Representative brightfield micrographs in Figure 6C show decreasing cell viability with increasing DOX concentration, indicated by reduced cell adherence and an increased prevalence of floating cells with spherical morphologies. High-throughput cell viability was quantified using a cell counter with Trypan Blue staining (see methods) (Fig. 6D). In conditions without DOX present, cell viability was comparable between unmodified and N-term variants at ND concentrations below 5 µM. The inclusion of DOX in ND formulations substantially reduced cell viability for both the unmodified and N-term variants in the sub-5 µM ND concentration range. The differences in efficacy for the various formulations were most pronounced at the highest ND concentration (∼50 μM), as summarized in Figure 6E. At this maximum concentration, N-term NDs elicited the largest reduction in cell viability, with effects comparable to the equivalent dose of free DOX. C-term NDs exhibited an intermediate cellular response relative to unmodified and N-term nanodiscs (Fig. S13). Together, these results demonstrate that R6W3-functionalization of NDs enhances DOX delivery to cells.

## DISCUSSION

In this work, we investigated how incorporation of the amphipathic CPP R6W3 into the MSP-based nanodiscs modulates membrane interactions, cellular uptake, and intracellular trafficking. Importantly, R6W3 functionalization did not disrupt nanodisc assembly, lipid organization, or protein-lipid interactions. All constructs formed monodisperse NDs with sizes and morphologies consistent with prior reports for DMPC nanodisc systems^12, 15, 37^, and exhibited biophysical signatures characteristic of properly assembled MSP-lipid complexes^38–40^. The absence of aggregation or rouleaux formation further supports that R6W3 fusion does not induce nanodisc self-association^41^. Together, these results confirm that functional differences observed across constructs arise from altered interfacial interactions rather than structural perturbations. Despite this structural equivalence, R6W3-functionalized NDs exhibited enhanced membrane association relative to unmodified NDs, with a strong dependence on peptide placement. N-terminal fusion consistently produced greater membrane binding than C-terminal fusion, indicating that peptide orientation governs interfacial engagement. Tryptophan residues are known to preferentially localize at the lipid headgroup-acyl chain interface and stabilize amphipathic helices^27, 42, 43^, and R6W3 adopts an amphipathic l1-helical secondary structure upon membrane binding^29^. These features suggest that N-terminal placement promotes more effective cooperative anchoring at the membrane surface. The observation of similar orientation-dependent trends in eGFP constructs further supports that this effect arises from peptide positioning rather than scaffold-specific interactions.

Consistent with enhanced membrane association, R6W3 functionalization increased cellular interaction and intracellular accumulation across all cell types examined. Notably, unmodified NDs possessed a baseline capacity for cell association, likely due to ApoA1-derived interactions with plasma membrane proteins, such as scavenger receptor class B type I (SR-B1) ^44–48^. Nonetheless, incorporation of R6W3 augmented this baseline interaction and enhanced cellular uptake. In agreement with prior studies demonstrating that CPP incorporation enhances nanoparticle uptake ^49, 50^, both N- and C-terminal R6W3 NDs exhibited increased intracellular fluorescence relative to unmodified NDs. Notably, intracellular fluorescence was quantified specifically within cytosolic regions, providing a more direct measure of internalization rather than whole-cell measurement.

Mechanistically, our data identifies CME as a dominant pathway for R6W3 ND internalization. Increased density of clathrin-coated pits, identified via AP2 labeling^51, 52^, together with intracellular colocalization of NDs with clathrin-coated vesicles, supports enhanced recruitment of CME machinery. Intracellular colocalization with clathrin-coated vesicles provides direct evidence of vesicular trafficking through CME. In contrast to prior CPP-functionalized ND systems that primarily report enhanced uptake^13–16^, these results directly establish a dominant endocytic route for peptide-functionalized NDs.

Early endosomes represent a major intracellular bottleneck for nanoparticle delivery, as internalized cargo is frequently retained within these compartments and prevented from accessing downstream intracellular targets. In this context, the reduced colocalization of R6W3-functionalized NDs with early endosomal markers suggests decreased sequestration within these compartments following clathrin-mediated uptake. This reduction in endosomal accumulation provides a plausible explanation for the enhanced intracellular delivery observed in our system. While the precise mechanism underlying this behavior cannot be definitively established from the present data, several possibilities may account for the observed decrease in early endosomal localization.

First, R6W3-functionalized NDs may exhibit enhanced endosomal escape, enabled by the amphipathic and membrane-active nature of the peptide. Second, R6W3 incorporation may promote accelerated trafficking through early endosomal compartments, reducing residence time and limiting accumulation. Third, R6W3-functionalized NDs may be diverted into alternative intracellular pathways that do not strongly overlap with early endosomal markers. Importantly, these mechanisms are not mutually exclusive, and the observed reduction in early endosomal accumulation likely reflects a combination of altered trafficking behaviors. Regardless of the specific mechanism, decreased sequestration within early endosomes is consistent with improved access to intracellular space.

Taken together, these results support a mechanistic model (Fig. 7) in which R6W3-functionalization enhances initial membrane association, promotes uptake through clathrin-coated pits and vesicles, and reduces accumulation within early endosomes following internalization. This sequence of events indicates that peptide-driven interfacial interactions not only govern cellular entry but also modulate intracellular trafficking by limiting endosomal sequestration.

**Figure 7:**
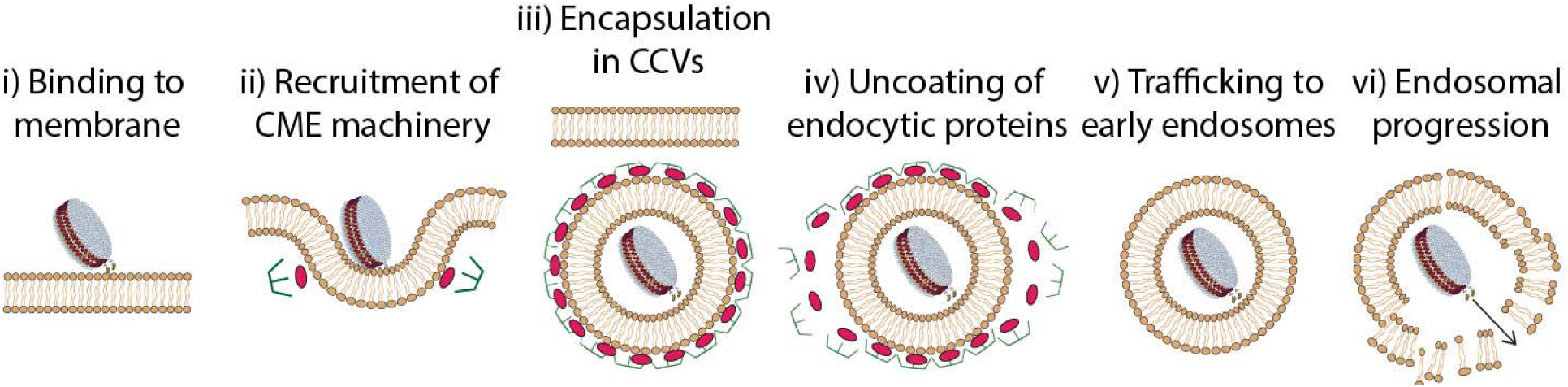
Proposed mechanism of cellular uptake and intracellular trafficking of R6W3-functionalized NDs.

These differences in intracellular trafficking translated directly to functional delivery outcomes. Despite relatively low DOX loading efficiency (∼2 DOX molecules per ND), R6W3-functionalized NDs produced greater reductions in cell viability than unmodified NDs at equivalent concentrations. This enhanced efficacy indicates that intracellular delivery efficiency, rather than total drug loading, is a dominant determinant of therapeutic performance. The limited DOX incorporation observed here is consistent with constrained partitioning of amphipathic DOX into tightly packed DMPC bilayers, as lipid composition is known to strongly influence drug loading efficiency^35, 53–56^. Nonetheless, improved cytotoxic response highlights the importance of intracellular trafficking in governing delivery outcomes.

## CONCLUSION

This work establishes that incorporation of an amphipathic CPP can modulate both the uptake pathway and intracellular fate of NDs without disrupting structural integrity. In contrast to prior approaches focused primarily on increasing cellular association, we show that peptide architecture and placement define endocytic routing and downstream trafficking behavior. The reduced early endosomal accumulation observed for R6W3-functionalized NDs highlights endosomal sequestration as a key barrier in ND-based delivery and suggests a strategy for overcoming this limitation. These findings provide a framework for engineering nanodisc-based delivery systems in which peptide design is used to control intracellular trafficking and improve functional delivery.

## MATERIALS AND METHODS

### Materials

1,2-dimyristoyl-sn-glycero-3-phosphocholine (DMPC), 2-dioleoyl-sn-glycero-3-phospho-L-serine (sodium salt) (DOPS), 1,2-dioleoyl-sn-glycero-3-phosphocholine (DOPC), 1,2-dipalmitoyl-sn-glycero-3-phosphoe-thanolamine-N-(cap biotinyl) (DPPE-Cap-Biotin) were purchased from Avanti Research. ATTO 647N-DPPE and ATTO 488 NHS Ester were purchased from ATTO-TEC (now available from Leica Microsystems). mPEG-Succinimidyl Valerate (mPEG-SVA) and Biotin-PEG-SVA were obtained from Laysan Bio, Inc. Tris(hydroxymethyl)aminomethane (Tris), phenylmethanesulfonylfluoride (PMSF), Poly-L-Lysine HCl (PLL), mineral oil, 1,6-Diphenyl-1,3,5-hexatriene (DPH), cOmplete™ Mini EDTA-free protease inhibitor tablets, 2- mercaptoethanol (BME), insulin from bovine pancreas, sucrose, and hydrocortisone were purchased from Sigma Aldrich. Janelia Fluor® 646 HaloTag® ligand (JF646) was obtained from Promega. Doxorubicin hydrochloride was purchased from Fisher Scientific 2-[4-(2-hydroxyethyl)piperazin-1-yl]-ethanesulfonic acid (HEPES), Texas Red™ 1,2-Dihexadecanoyl-*sn*-Glycero-3-Phosphoethanolamine Triethylammonium Salt (Texas Red-DHPE), Fetal Bovine Serum (FBS), Triton X-100, CellMask™ Deep Red plasma membrane stain, CellLight™ Early Endosomes-RFP, trypan blue solution (0.4%), imidazole, and NeutrAvidin protein were purchased from Thermo Fisher Scientific. Isopropyl-beta-D-thiogalactoside (IPTG) was obtained from Gold Biotechnology. Trypsin (0.05%), penicillin/streptomycin/L-glutamine (PSLG), F12 HAM’S nutrient mixture, and DMEM high glucose without L-glutamine, and sodium pyruvate, and phenol red free DMEM were purchased from Cytiva Life Sciences.

### Expression and Purification of Proteins

The his-tagged MSP1D1 construct (pMSP1D1) was a gift from Stephen Sligar (Addgene plasmid # 20061). Mutant MSP1D1 constructs containing R6W3 at either the N- or C- terminus were generated using plasmid editing services from Genscript, where nucleotides coding for R6W3 and the corresponding linker were inserted into pMSP1D1 plasmids. MSP1D1 constructs containing R6W3 were expressed in the original pET28a vector backbone, while unmodified MSP1D1 was subcloned into the pRSET high-copy expression vector. All plasmids were propagated in *Escherichia Coli* (*E. coli*) DH5α, and recombinant protein expression was carried out in *E. coli* BL21 cells. Protein expression was performed following a previously established protocol for MSP1D1, with modification ^57^. Transformed BL21 cultures were grown in 2XTY media at 37°C with shaking at 220 rpm until cells reached an OD600 of ∼0.6. The incubation temperature was reduced to 30°C and protein expression was induced with 1mM of IPTG. The cells were incubated for an additional 3 hours following induction and subsequently harvested by centrifugation at 9,000 g for 20 mins. Cell pellets were resuspended in 1% (v/v) Triton X-100 and lysis buffer containing 500 mM Tris-HCl (pH 8.0), 5% (v/v) glycerol, 10 mM BME 1 mM PMSF, cOmplete™ protease inhibitor tablets. Cells were lysed by tip sonication (on ice) at 60% amplitude for four cycles of 2 mins on and 2 mins off. The lysate was clarified by using ultracentrifugation at 40,000 rpm (103,800 g, s50-A rotor) for 40 mins at 4 °C, where the supernatant was collected for further purification. His-tagged proteins were purified using Ni-NTA affinity chromatography. Ni-NTA agarose resin (6 mL of 50% slurry, 3 mL bed volume) was pre-equilibrated with buffer containing 400 mM Tris-HCl (pH 8.0), 10 mM BME, and 1 mM PMSF. The lysate was incubated with the resin for 1 hour at 4 °C prior to being loaded to the column. Unbound proteins were rinsed away with Tris-HCl buffer (pH 8.0) containing increasing concentrations of imidazole and 1M NaCl. His-tagged proteins were eluted with Tris-HCl (pH 8.0) containing 200 mM imidazole. Eluted fractions were pooled and dialyzed for 24 hours using a Slide-A-Lyzer dialysis cartridge against HEPES buffer (25 mM HEPES, 150 mM NaCl, pH 7.4) to remove imidazole and exchange buffers. The pRSET-his-eGFP construct was a gift from Jeanne Stachowiak (Addgene plasmid #113551). Mutant eGFP constructs containing R6W3 at either the N- or C- terminus were generated by Genscript using the pRSET-eGFP construct as a template. Each plasmid was amplified in *E. Coli* DH5α, and then protein expression was carried out using *E. coli* BL21 cells. Protein expression and purification was performed following similar methods described for membrane scaffold proteins with slight modifications. Once transformed BL21 cells reached an OD600 of ∼0.6, the incubation temperature was reduced to 18°C, protein expression was induced with 1mM of IPTG and allowed to incubate for 16 hours prior to being harvest by centrifugation at 9,000 g for 20 mins.

All protein concentrations were determined by UV-Vis spectroscopy using extinction coefficients (280 nm) of 21,430 M^-1^ cm^-1^ for unmodified MSP1D1, 37,930 M^-1^ cm^-1^ for R6W3-functionalized MSP1D1 mutants, and an extinction coefficient (488 nm) of 56,000 M^-1^ cm^-1^ for all eGFP constructs. Protein purity and molecular weight were assessed by SDS-Page electrophoresis.

### Synthesis of Nanodiscs

MSP1D1 and mutant nanodiscs were assembled using a previously established detergent-based reconstitution method^10^. Briefly, DMPC lipid aliquots stored at -80°C were thawed, and organic solvents were removed under a stream of nitrogen, followed by vacuum desiccation for 2 hours to form dry lipid films. The lipid films were resolubilized in HEPES buffer containing 40 mM sodium cholate, combined with protein at a lipid:protein molar ratio of 150:1, and placed on a nutating mixer for 30 minutes. The mixture was subsequently transferred to a 10 kDa MWCO dialysis cartridge for detergent removal. The resulting nanodisc mixture was further purified by size-exclusion chromatography using a Superdex 200 Increase 10/300 GL column (Cytiva) on an AKTA pure system (Cytiva). Fractions corresponding to nanodisc-containing elution peaks were collected, pooled and concentrated using a 50 kDa MWCO centrifugal filter (Millipore Sigma).

Nanodiscs containing Texas Red-DHPE were also assembled using the same detergent based protocol described above with the following modifications. Briefly, starting lipid aliquots were combined to yield a 98% DMPC and 2% Texas Red-DHPE molar percentage, then organic solvents were evaporated to form dry lipids films. Size exclusion chromatography was replaced with affinity chromatography using Ni-NTA agarose beads. Following dialysis, the nanodisc mixture was incubated with Ni-NTA beads for 1 hour at 4°C. Bound nanodiscs were eluted using 300 mM imidazole, recovered, and buffer exchanged prior to being concentrated by ultracentrifugation. Nanodiscs containing DOX were assembled using the same detergent based reconstitution protocol described above, with the following modification. DOX was resuspended in water and incubated with detergent solubilized DMPC lipid and MSP1D1 (or mutant protein) at a lipid:protein:DOX molar ratio of 150:1:10 prior to being dialyzed. Following dialysis, DOX-loaded NDs were purified using repeated dilution and concentration cycles (3 cycles, approximately 8-fold dilution each cycle) in a 50 kDa MWCO centrifugal filter, rather than size-exclusion chromatography.

For protein-dye labeling, ATTO 488 NHS-Ester (ATTO Tec) was dissolved in dimethyl-sulfoxide and mixed with MSP1D1 constructs in HEPES buffer followed by an incubation of 30 minutes in the dark at room temperature. Unreacted dye was removed from solution using a 7 kDa MWCO spin desalting column (ThermoFisher).

Dyes and drug concentrations were determined by UV-Vis spectroscopy using a NanoDrop spectrophotometer. Protein concentrations were determined using absorbance values at 280 nm, applying extinction coefficients of 21,430 M^-1^ cm^-1^ for MSP1D1 and 37,930 M^-1^ cm^-1^ for R6W3-containing MSP1D1 constructs. Texas-Red concentration was determined using absorbance values at 595 nm, applying an empirically determined extinction coefficient of 65,575 M^-1^ cm^-1^,with an empirically determined correction factor of 0.29 for the 280 nm/595 nm absorption ratio. ATTO 488 dye concentration was determined using absorbance values at 488 nm, applying an extinction coefficient of 90,000 M^-1^ cm^-1^ with a correction factor of 0.09 for the 280 nm/488 nm absorption ratio. DOX concentrations were determined using absorbance values at 480 nm, applying an empirically determined extinction coefficient (i.e., from a standard curve) of 8,312 M^-1^ cm^-1^, with an empirically determined correction factor of 0.82 for the 280 nm/480 nm absorption ratio.

### Dynamic Light Scattering

Hydrodynamic diameter measurements were determined in HEPES buffer by dynamic light scattering (Nano Particle Analyzer SZ-100, Horiba Scientific).

### Transmission Electron Microscopy

Negative-stain transmission electron microscopy (TEM) micrographs were obtained on a ThermoFisher Talos F200C G1 microscope using an operating voltage of 80 kV and a zoom of 92,000x. All formulations of nanodiscs were diluted to 1 μM in HEPES buffer. Formvar/carbon-coated nickel grids (200 mesh) were treated in a UV/ozone chamber for 4 minutes. 4 μL of sample was applied to each grid and blotted dry using filter paper. The samples were then stained twice with 4 μL of 1% uranyl acetate (UA). Size measurements of nanodiscs were quantified using ImageJ.

### Fluorescence Anisotropy

Tryptophan fluorescence anisotropy was measured using a spectrofluorometer (JASCO FP-8500). Samples containing 1 μM protein in methacrylate cuvettes were excited at 280 nm and fluorescence anisotropy was measured using a with detection range 300-480 nm at room temperature. Anisotropy values (r) were calculated from polarized emission intensities as shown in eq 1^34^.

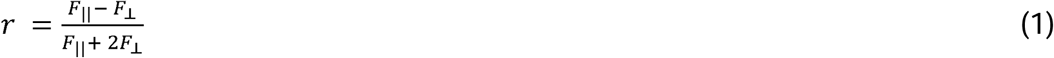

Temperature-dependent DPH fluorescence anisotropy measurements were obtained using the same JASCO spectrofluorometer. Small unilamellar vesicles (SUVs, 60 μM total lipid) composed of DMPC were prepared by probe tip sonication. Debris from tip sonication was removed by centrifugation prior to use. DPH was added to SUV- and ND- containing solutions at 100:1 lipid:DPH molar ratio. Samples were excited at 355 nm and emission intensities were monitored at 425 nm, as the temperature increased gradually from 10 to 40°C. Temperature-dependent fluorescence anisotropy curves were fit using the method of least-squares regression to an empirical sigmoidal phase transition model eq2^58^. The fitting parameters *r_max_*, *r_min_*, *T_m_*, and n correspond to the maximum anisotropy, minimum anisotropy, phase transition temperature and cooperativity index, respectively.

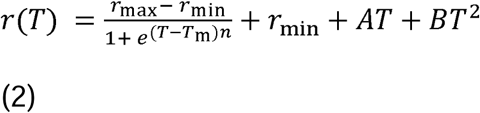

### Epifluorescence Microscopy

Imaging was performed on a fluorescence microscope (ECHO Revolve) using transmitted light and fluorescence imaging. Transmitted light images were acquired using phase contrast optics with an exposure time of 55 ms. Fluorescence imaging was performed using a FITC filter cube with an exposure time of 670 ms. The low camera gain setting was used, and light intensity was set to 50% for all images. An Olympus LucPlanFLN 20x/0.45 NA phase contrast objective was used for image acquisition. Phase contrast optics were used for transmitted light imaging, while the optical configuration was switched to brightfield when acquiring fluorescent images.

### Confocal Microscopy

Confocal imaging was performed on a laser scanning confocal microscope (STELLARIS 5, Leica microsystems). Images were acquired using a Leica HC PL APO 63x/1.4 NA oil-immersion objective. Image settings were set to the following unless explicitly stated otherwise: pixel size of 70 nm, pinhole size of 1.0 Airy Units, and scan speed to 400 Hz.

### Imaging of Nanodiscs

NDs were imaged by adsorbing them to untreated glass coverslips (No. 1.5 glass) at a concentration of 25 pM. Silicone gaskets were used to create imaging wells (Grace BioLabs). Samples were allowed to adhere to glass surfaces via electrostatic interactions, after which wells were gently rinsed with HEPES buffer to remove excess NDs in solution. Confocal microscopy was used for image acquisition with the following settings. ATTO 488 and Texas Red were excited using lasers at 488 and 561, respectively. Fluorescence emission was collected over the wavelength ranges of 493-550 nm and 583-734 nm for ATTO 488 and Texas Red, respectively.

### Preparation of Giant Unilamellar Vesicles (GUVs)

Giant unilamellar vesicles (GUVs) were synthesized using an inverted emulsion method^59^. A lipid solution containing DOPS:DOPC:ATTO 647N DHPE:DPPE-Cap-Biotin at a 49:49:1:1 molar ratio was dried under nitrogen and vacuum desiccated for 2 hours. The dried lipid film was resuspended in 1 mL of mineral oil to reach a total lipid concentration of 500 μM. The lipid solution was mixed, incubated at 50°C for 20 mins and layered (300 μL) over 400 μL of HEPES buffer. Samples were stored vertically in the dark for 45 min to form a lipid monolayer at the oil-aqueous interface. An inverted emulsion was formed by mixing 20 μL of sucrose buffer (300 mM Sucrose, 25 mM HEPES, pH 7.4) with 600 μL of the lipid solution in mineral oil. The inverted emulsion was pipetted above the lipid monolayer and centrifuged at 2,500 x g for 10 mins for GUV pellets at the bottom of the centrifuge tube. The supernatant was aspirated using a glass Pasteur pipet to allow 100 μL of GUV- containing solution to be resuspended in HEPES buffer.

### Tethered Vesicle Assay

The vesicles prepared as described in **Preparation of Giant unilamellar vesicles (GUVs)** were tethered to glass clover slips using a previously established protocol^60^. Briefly, No. 1.5 glass coverslips and silicon gaskets were cleaned with heated Hellmanex solutions and allowed to dry under nitrogen. Gaskets were affixed to the coverslips, followed by a 20-min incubation of each well with a mixture of PLL-PEG:PLL-PEG-biotin (49:1 molar ratio) to passivate the glass surface. Wells were rinsed with HEPES buffer to remove excess passivation agents. Neutravidin was incubated to reach a concentration of 0.2 mg/mL for a total of 10 minutes then rinsed with HEPES buffer. Each well was incubated with vesicles at a lipid concentration of 2.5 μM and rinsed with HEPES buffer. Lastly, wells were then incubated with increasing concentrations of Texas-Red- and ATTO 488- labeled nanodiscs and imaged using confocal microscopy. All vesicles were imaged at their equatorial plane. GUVs incubated with nanodiscs were excited with lasers at 488, 561, and 638 nm. The fluorescence emission detection wavelengths were set to 500-550 nm for ATTO488, 565-625 nm for Texas Red, and 645-749 nm for ATTO 647N. GUVs incubated with eGFP were excited with excitation lasers at 488, and 638 nm. The fluorescence emission detection wavelengths were set to 493-593 nm for eGFP and 643-834 nm for ATTO 647N.

### Quantification of GUV Associated Fluorescence

GUV associated fluorescence was quantified using a previously developed, MATLAB-based vesicle analysis algorithm ^61^. In short, the algorithm identified the center of individual vesicles imaged and generated radial fluorescent intensity profiles as a function of distance from the center of the vesicle for each fluorescent channel. Membrane fluorescence was quantified by fitting the intensity profile to a gaussian function. Specifically, the gaussian amplitudes corresponded to fluorescence intensity on the surface of the GUVs. The peaks obtained were then used to perform a ratiometric analysis between different fluorescent channels to quantify the relative signal on the membrane surface. For all fluorescence-based measurements, intensity values were normalized by their corresponding labeling ratios (i.e., ATTO 488 dyes per protein or number of lipids per ND as determined by Texas-Red fluorescence) to account for differences in protein labeling efficiency and lipid incorporation in nanodiscs.

### Cell Culture

SUM159 cells genetically engineered to express a Halo-Tag fused to both alleles of the AP-2 c2 subunit was a gift from the Kirchhausen laboratory at Harvard University^62^. SUM159 cells were maintained in a 1:1 mixture of Ham’s F-12 nutrient mix and DMEM high glucose. The growth media was supplemented with fetal bovine serum (FBS, 5%), HEPES (10 mM), insulin (5 μg/mL), hydrocortisone (1 μg/mL), and PSLG (1%). The pH of the media was adjusted to 7.4 using filtered NaOH (5 M stock solution). RPE and HeLa cells were gifts from the Stachowiak Laboratory at the University of Texas at Austin^63, 64^. RPE cells were grown in a 1:1 mixture of Ham’s F-12 nutrient mix and DMEM high glucose supplemented with 10% FBS, 1% PSLG, and 20 mM of HEPES. HeLa cells were grown in DMEM supplemented with 10% FBS, and 1% PSLG. All cells were maintained in a humidified incubator set to 37°C with 5% CO_2_.

### Nanodisc-Cell Association Studies

To assess broad nanodisc interactions with cells, ATTO 488 labeled NDs were incubated with either SUM159, HeLa, or RPE cells and imaged using epifluorescence microscopy. Cells were seeded at a density of 3 x10^4^ cells/mL in 24 well plates (Corning Inc.) 16 hours prior to imaging and incubated with NDs for 1 hour at 37°C with 5% CO_2_. Following incubation, wells were rinsed with PBS and replenished with phenol red free growth media. Cells were imaged using the epifluorescence microscopy settings described above, with FITC excitation for ATTO488 and transmitted light for cell surface visualization.

ATTO 488 fluorescence associated with the cell was quantified using a custom image analysis method. Cells were segmented from the transmitted light images using the open-source software Cellpose^65^, and the resulting masks were imported into ImageJ and MATLAB for analysis. Separate masks corresponding to cell-associated and background regions were generated and applied to the ATTO 488 fluorescence images to quantify fluorescence inside and outside the cell. To account for regions within segmented cells that contained little to no ATTO 488 signal, a threshold based weighting factor approach was used. Cell associated ATTO488 signal was defined as the mean intensity (〈*I_cell_*〉) exceeding three standard deviations (σ*_bg_*) above the background mean intensity (〈*I_bg_*〉). Background fluorescence was obtained by measuring the mean intensity outside segmented cells. The cell associated fluorescence was calculated by taking the product shown in equation 3, where the weighting factor is defined as the number of cell-associated pixels above background noise (*N*_〈*I_cell_*〉_ > 〈*I_bg_*〉 +3σ*_bg_* ) divided by total cell-associated pixels (*N*_<_*I_cell_*_>_). A factor of 10_ was chosen to re-scale cell associated fluorescence values.

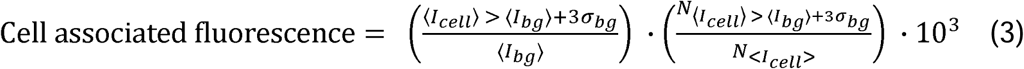

This image analysis and quantification procedure were applied identically across all experimental conditions.

### Nanodisc Internalization Studies

To assess cellular internalization of NDs, ATTO 488 labeled NDs were incubated with SUM159, HeLa, or RPE cells and imaged using confocal microscopy. Cells were seeded at a density of 1x10^5^ cells/mL onto 35 mm glass bottom dishes (CellVis) 16 hours prior to imaging. Cells were rinsed with PBS to remove any cellular debris then incubated with ATTO 488 labeled NDs for 1 hour at 37°C in a humidified incubator with 5% CO_2_. Following ND incubation, cells were stained with a plasma membrane dye, CellMask™ Deep Red by incubation in phenol red free containing media with dye diluted 1:1000x from the commercial stock for 15 mins at 37°C. Cells were rinsed with PBS and replenished with phenol red free growth media for imaging. Cells were imaged with the same confocal microscope mentioned above using 488 nm and 638 excitation lasers. Fluorescence emission was collected over wavelength ranges of 493 – 600 nm and 643 – 800 nm, for ATTO 488 and CellMask™, respectively. For single cell imaging, the zoom factor adjusted such that a square pixel size was 56.72 nm, and images were acquired at a scan speed of 600 Hz.

Intracellular nanodisc signal was quantified using confocal images of single cells. The cytosolic region of each cell was segmented using Cellpose based on the CellMask™ fluorescent channel. The masks were applied to the ATTO 488 channel to analyze the mean intracellular fluorescence. Internal ATTO 488 fluorescence was quantified using a similar threshold-based, weighting factor as described above (eq.4). Here, fluorescence intensities were not normalized by background fluorescence.

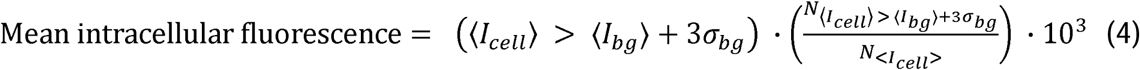

### Internalization Tracking of Nanodiscs

To examine the internalization pathway of nanodiscs, AP-2 was fluorescently labeled within the SUM159 cells using a HaloTag Ligand. SUM159 cells were seeded at a density of 1x10^5^ cells/mL onto 35 mm glass bottom dishes 16 hours prior to imaging. To visualize HaloTag-labeled AP-2, SUM159 cells were incubated with a membrane permeable HaloTag ligand conjugated to JF646 at a final concentration of 100 nM for 15 minutes before imaging. Excess ligand was removed by rinsing with phenol red-free culture medium, which also served as the imaging buffer. SUM159 cells were imaged using confocal microscopy with the following image acquisition parameters: ATTO 488 and JF646 were excited using 488 nm and 638 nm lasers, respectively. Fluorescence emission was collected over wavelength ranges 493 – 580 nm for ATTO 488 and 643-750 nm for JF646. Z-stack images were acquired with a step size of 0.05 μm per slice. The pinhole was set to 0.5 and the scan speed was set to 600 Hz.

To quantify the endocytic pits at the plasma membrane, the density of AP-2-labled pits was measured. Single cell images were segmented using Cellpose, and the resulting masks were imported to imageJ for analysis. The number of puncta visualized in the AP-2-JF646 fluorescence channel along with the corresponding cell area within the region of interest were quantified using ImageJ. Clathrin-coated pit density was calculated as the number of puncta divided by the cell area. Nanodiscs adsorption to the plasma membrane was quantified by measuring the mean ATTO 488 fluorescence intensity at the cell surface.

To investigate endosomal trafficking, endosomal compartments were fluorescently labeled using CellLight™ reagents. SUM159 cells were seeded at a density of 1x10^5^ cells/mL in a 35 mm glass bottom dish. CellLight™ Early Endosome-RFP (Rab5) reagent was added at the time of seeding using an excess particles per cell ratio, following the manufacturer’s recommendation. Cells were incubated for 16 hours to allow the expression of RFP-tagged Rab5. Following expression, cells were rinsed with PBS and incubated with ATTO 488 labeled nanodiscs for 1 hour at 37°C. Cells were then imaged using confocal microscopy. ATTO 488 and RFP were excited using 488 nm and 561 nm lasers, respectively. The pinhole was adjusted to 0.5 and the scan speed was increased to 600 Hz.

### Cell Viability

Cell viability was quantified using a TC20™ automated cell counter (BioRad) with disposable counting slides. For sample preparation, SUM159 cells were seeded in 24 well plates at a density of 3 x10^4^ cells/mL and allowed to adhere for 16 hours at 37°C. Wells were rinsed with PBS to remove any cell debris and incubated with either DOX-containing or DOX-free samples in phenol red free growth media for 24 hours. Culture media within the wells was then aspirated and collected in centrifuge tubes. Remaining cells were detached by trypsinization and were also pooled into the centrifuge tubes. Cells then underwent pelleting by centrifugation at 300 x g for 5 min. Cell pellets were resuspended in fresh growth media and samples were mixed at a 1:1 ratio with 0.4% (w/v) trypan blue.

## ACKNOWLEDGEMENTS

This work was supported in part by the National Institutes of Health through R35GM147333 to B.S. Pizarro and W. F. Zeno.

