## Supporting Information for "Amphiphilic Peptide Fusion Promotes Endocytic Uptake of Nanodiscs"

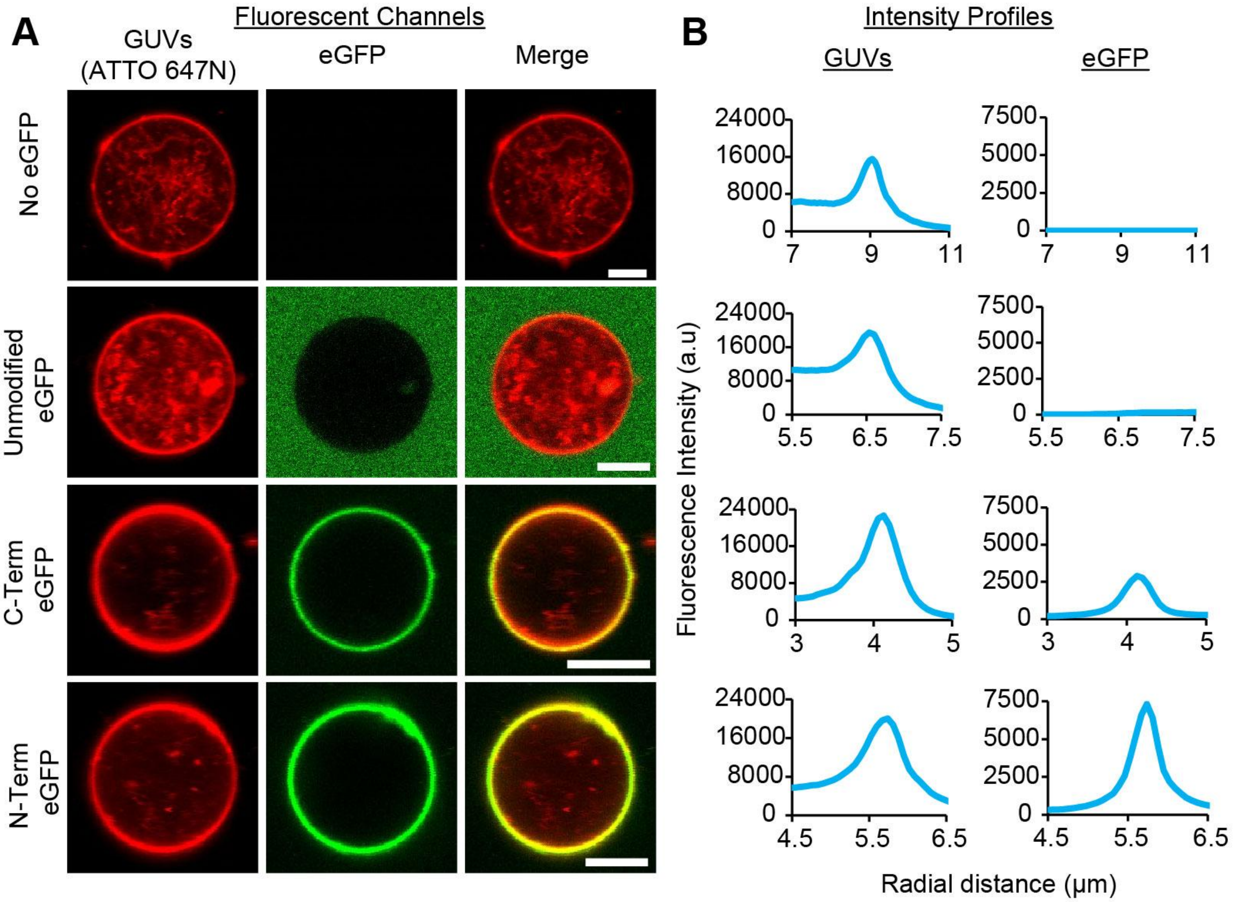


**Figure S1: R6W3-functionalized eGFP associates with model membranes.** (A) Representative confocal fluorescence images of GUVs (ATTO 647N) incubated with and without eGFP and its R6W3-containing variants. (B) Radial fluorescence intensity profiles of lipid and eGFP channels. Scale bar: 5 µm.

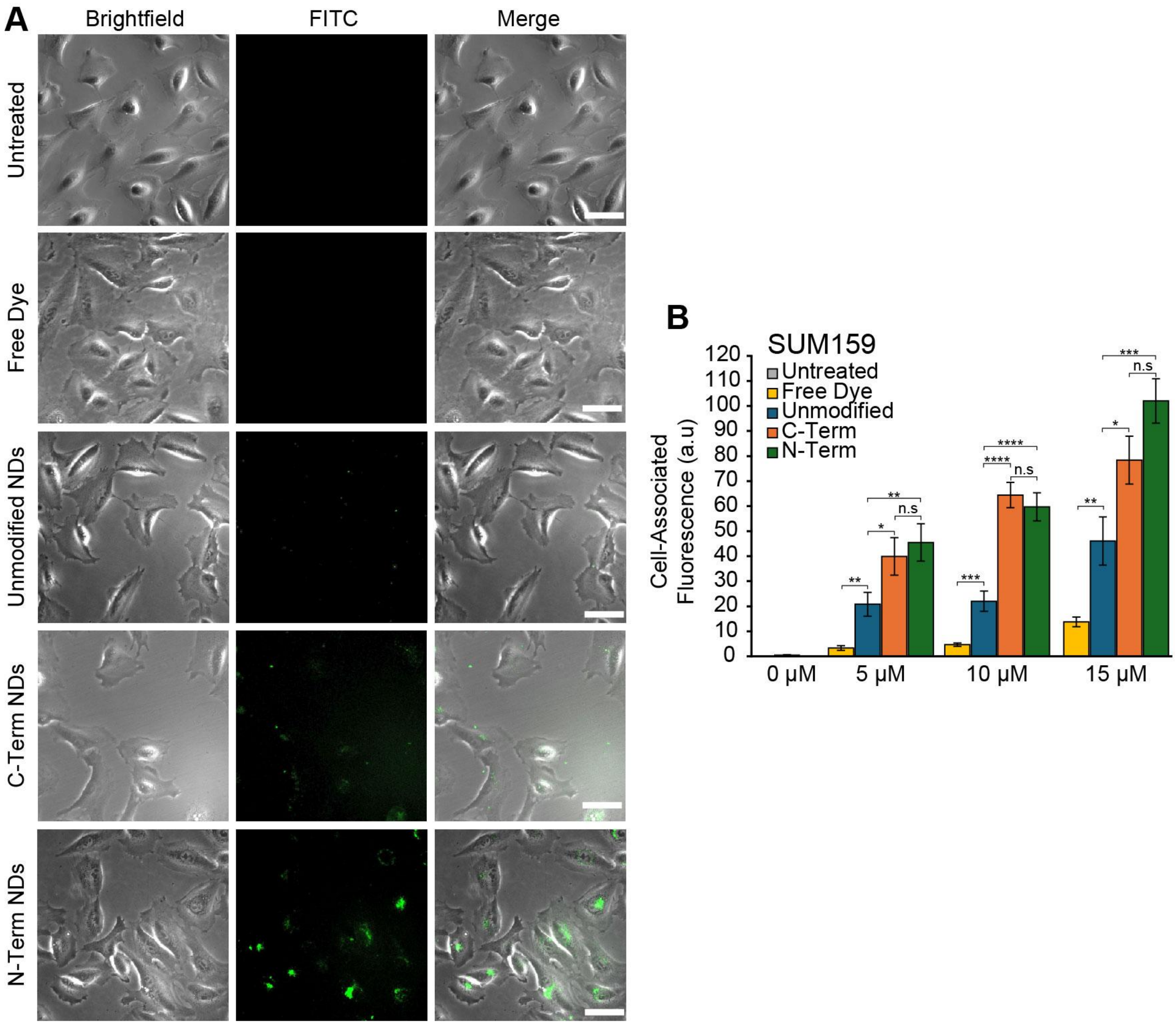


**Figure S2: R6W3-functionalized NDs enhances cellular association in SUM159 cells.** (A) Representative brightfield and fluorescence images of SUM159 cells following incubation with or without fluorescently labeled species. (B) Quantification of cell-associated fluorescence in SUM159 cells as a function of ND or free dye concentration. Data are presented as mean $\pm$ SEM. Statistical significance was determined by two tailed unpaired Student's t-test (* = p  < .05, ** = p < .01, *** = p < .001, **** = p < .0001, n.s = not significant). Scale bars: 50 µm.

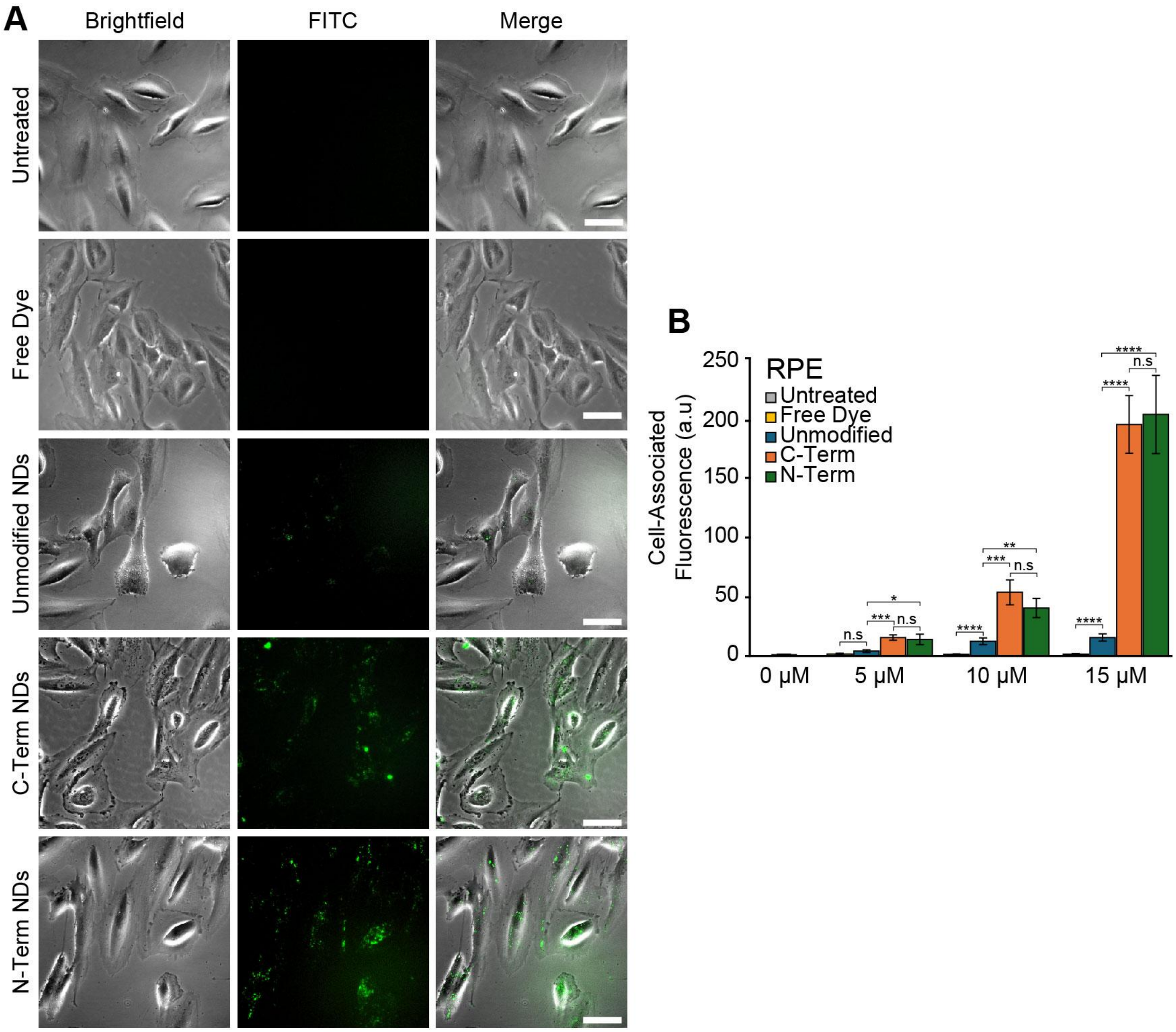


**Figure S3: R6W3-functionalized NDs enhances cellular association in RPE.** (A) Representative brightfield and fluorescence images of RPE cells following incubation with or without fluorescently labeled species. (B) Quantification of cell-associated fluorescence in RPE cells as a function of ND or free dye concentration. Data are presented as mean ± SEM. Statistical significance was determined by two tailed unpaired Student's t-test (* = p < .05, ** = p < .01, *** = p < .001, **** = p < .0001, n.s = not significant). Scale bars: 50 µm.

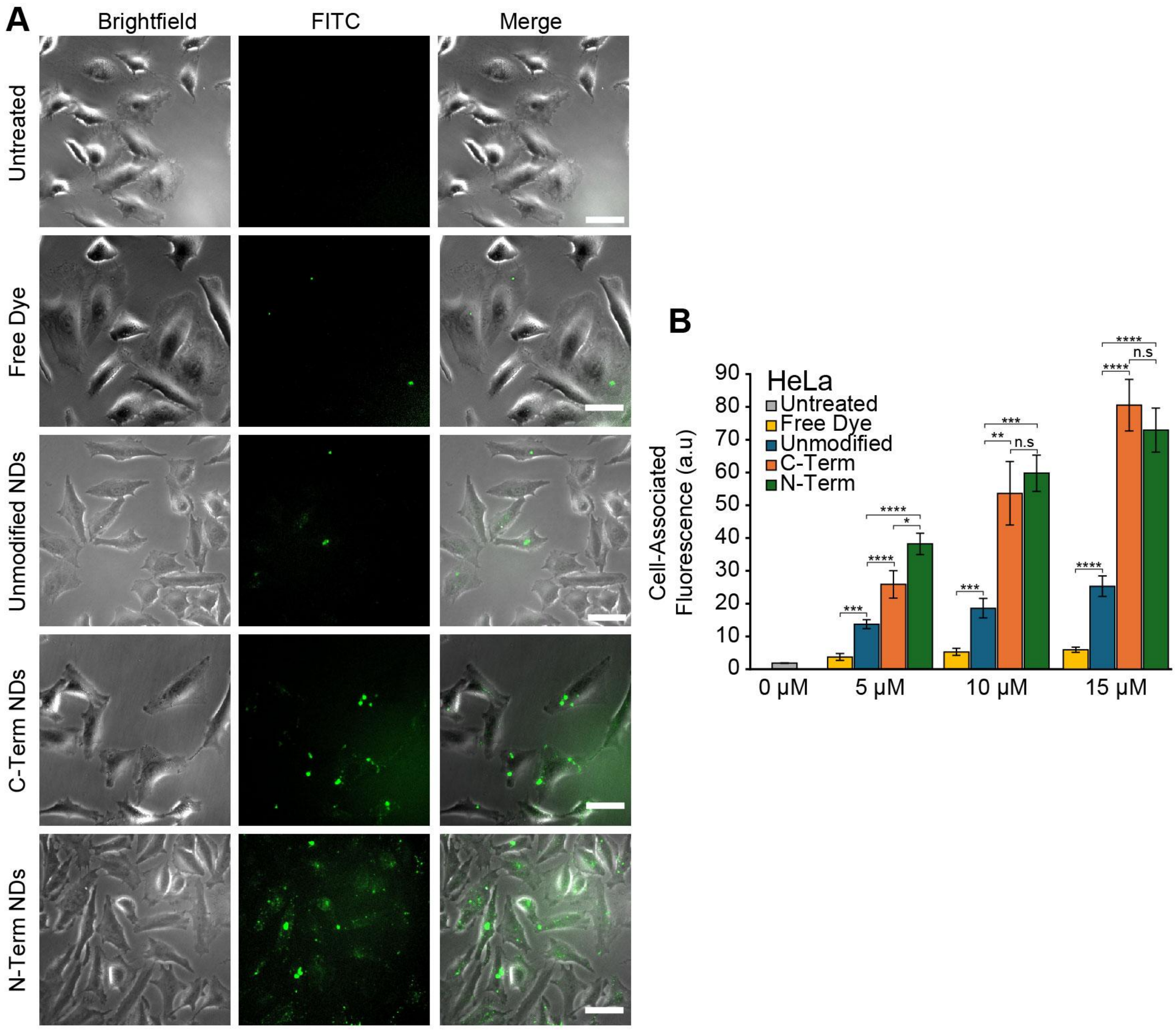


**Figure S4: R6W3-functionalized NDs enhances cellular association in HeLa.** (A) Representative brightfield and fluorescence images of HeLa cells following incubation with or without fluorescently labeled species. (B) Quantification of cell-associated fluorescence in HeLa cells as a function of ND or free dye concentration. Data are presented as mean ± SEM. Statistical significance was determined by two tailed unpaired Student's t-test (* = p < .05, ** = p < .01, *** = p < .001, **** = p < .0001, n.s = not significant). Scale bars: 50 µm.

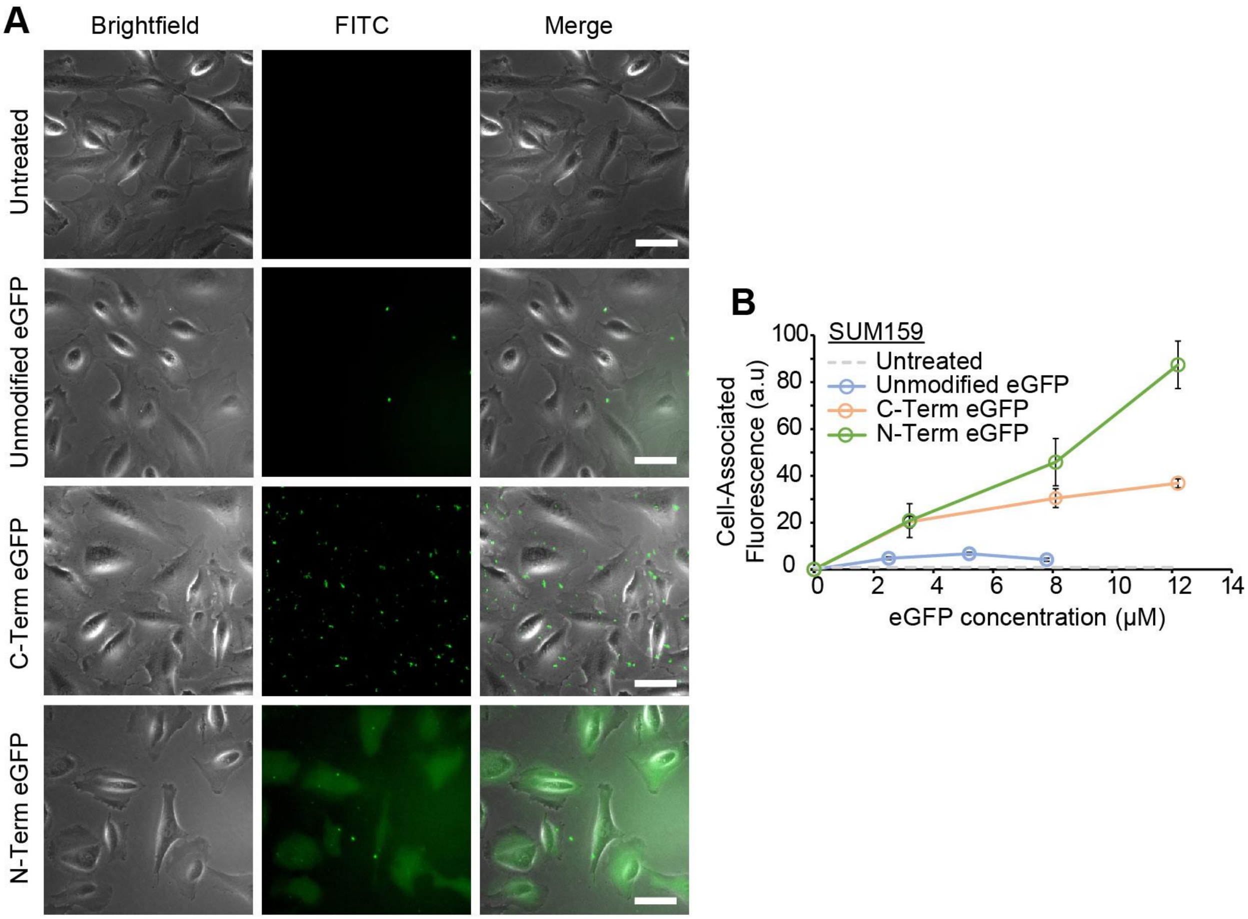


**Figure S5: R6W3-functionalized eGFP enhances cellular association in SUM159 cells.** (A) Representative brightfield and fluorescence images of SUM159 cells following incubation with unmodified, C-term, or N-term eGFP or untreated controls. (B) Quantification of cell-associated fluorescence in SUM159 cells as a function of eGFP concentration. Data are presented as mean ± SEM. Scale bars: 50 µm.

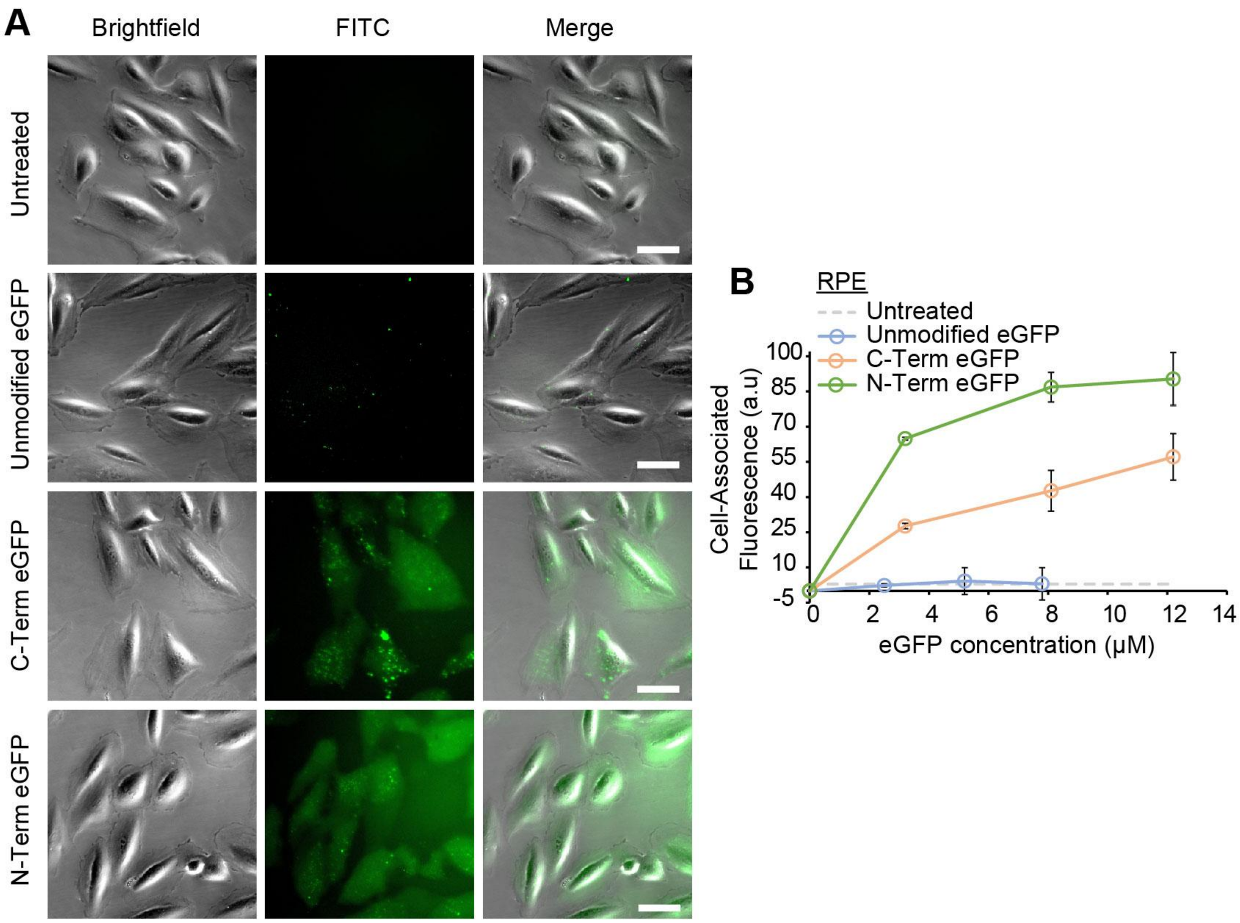


**Figure S6: R6W3-functionalized eGFP enhances cellular association in RPE cells.** (A) Representative brightfield and fluorescence images of RPE cells following incubation with unmodified, C-term, or N-term eGFP or untreated controls. (B) Quantification of cell-associated fluorescence in RPE cells as a function of eGFP concentration. Data are presented as mean ± SEM. Scale bars: 50 µm.

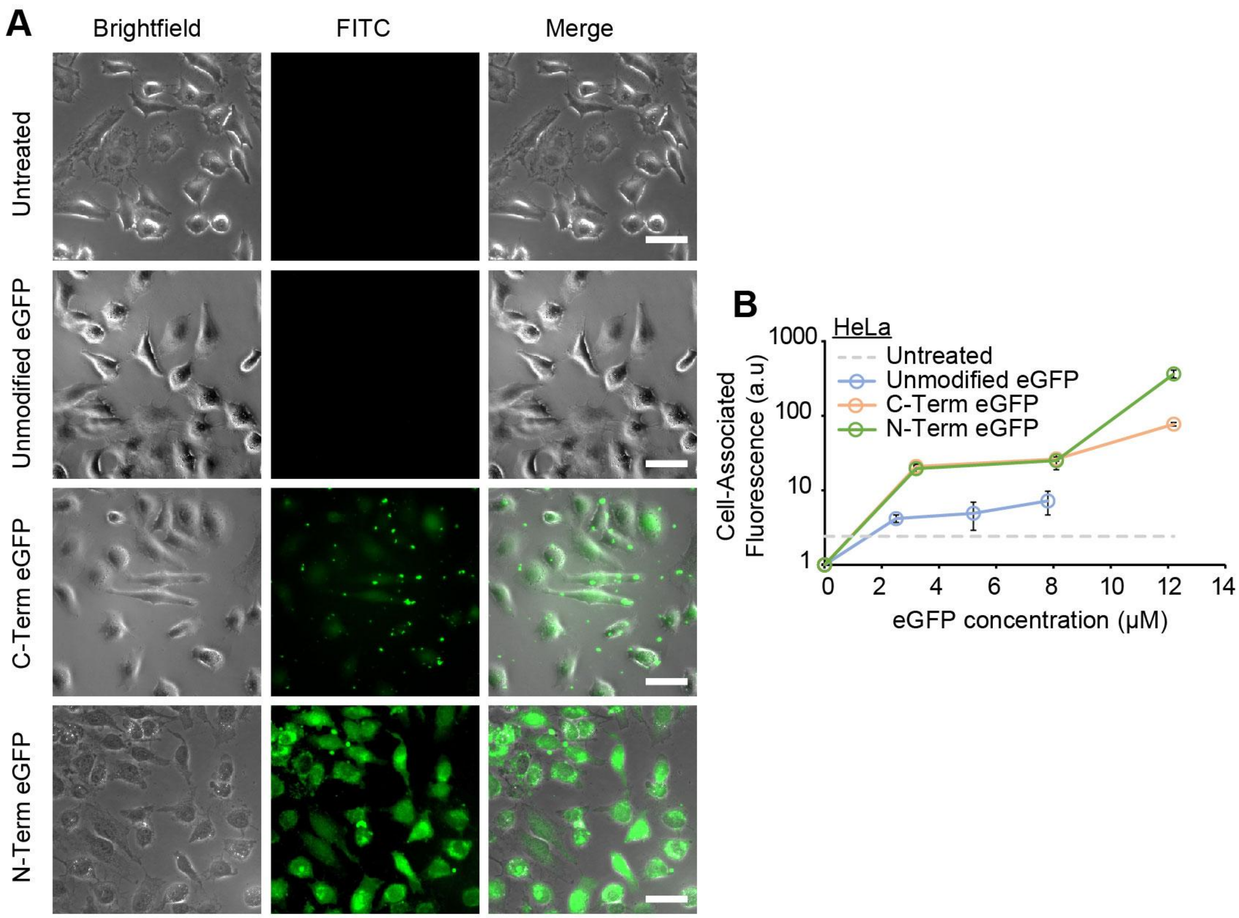


**Figure S7: R6W3-functionalized eGFP enhances cellular association in HeLa cells.** (A) Representative brightfield and fluorescence images of HeLa cells following incubation with unmodified, C-term, or N-term eGFP or untreated controls. (B) Quantification of cell-associated fluorescence in HeLa cells as a function of eGFP concentration. Data are presented as mean ± SEM. Scale bars: 50 µm.

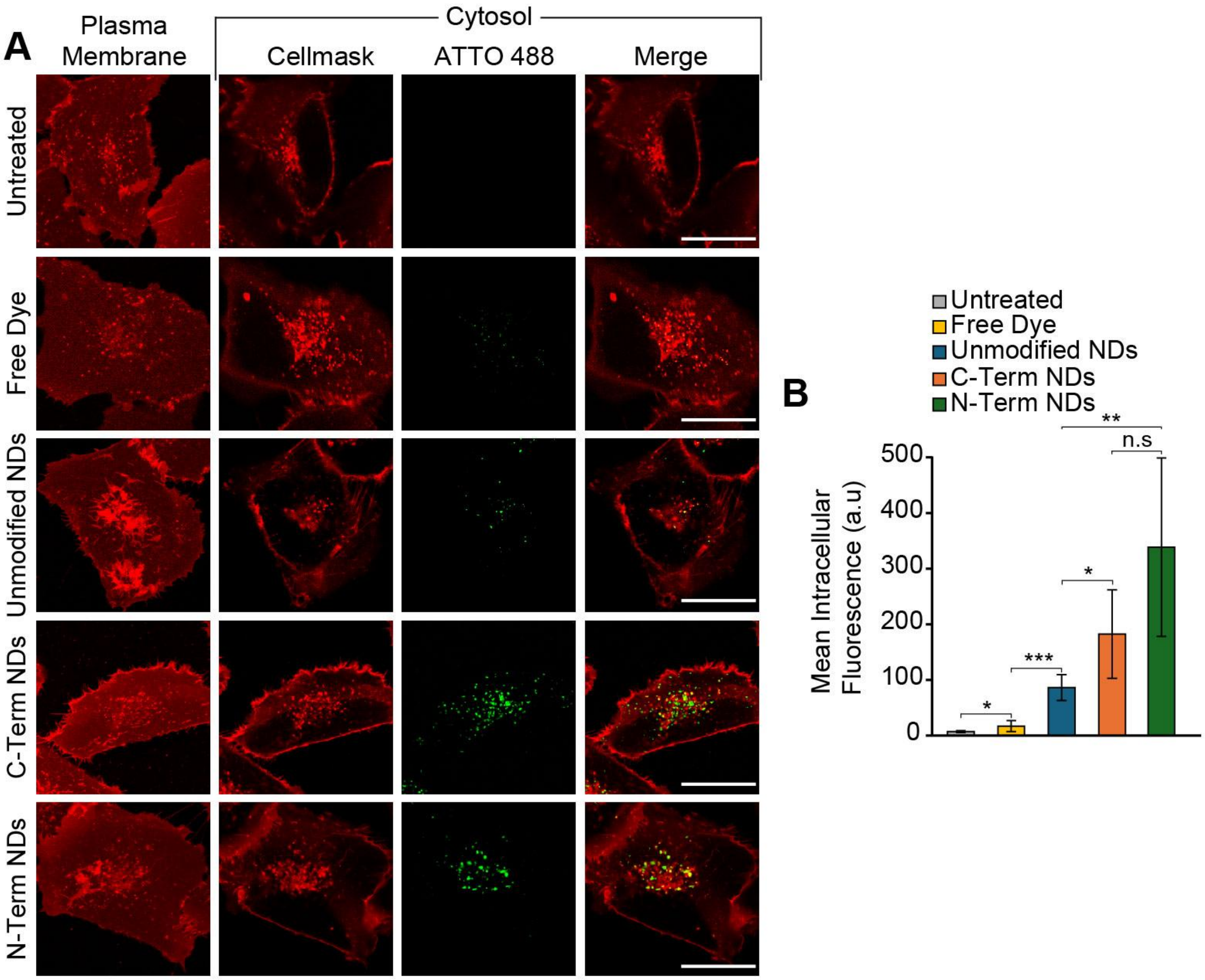


**Figure S8: R6W3-functionalized NDs undergo cellular internalization in SUM159 cells.** (A) Representative confocal images of SUM159 cells incubated with fluorescent NDs. (B) Quantification of intracellular fluorescence in SUM159 cells. Data are presented as mean ± SD. Statistical significance was determined by two tailed unpaired Student's t-test (* = p < .05, ** = p < .01, *** = p < .001, **** = p < .0001, n.s = not significant). Scale bars: 25 μm.

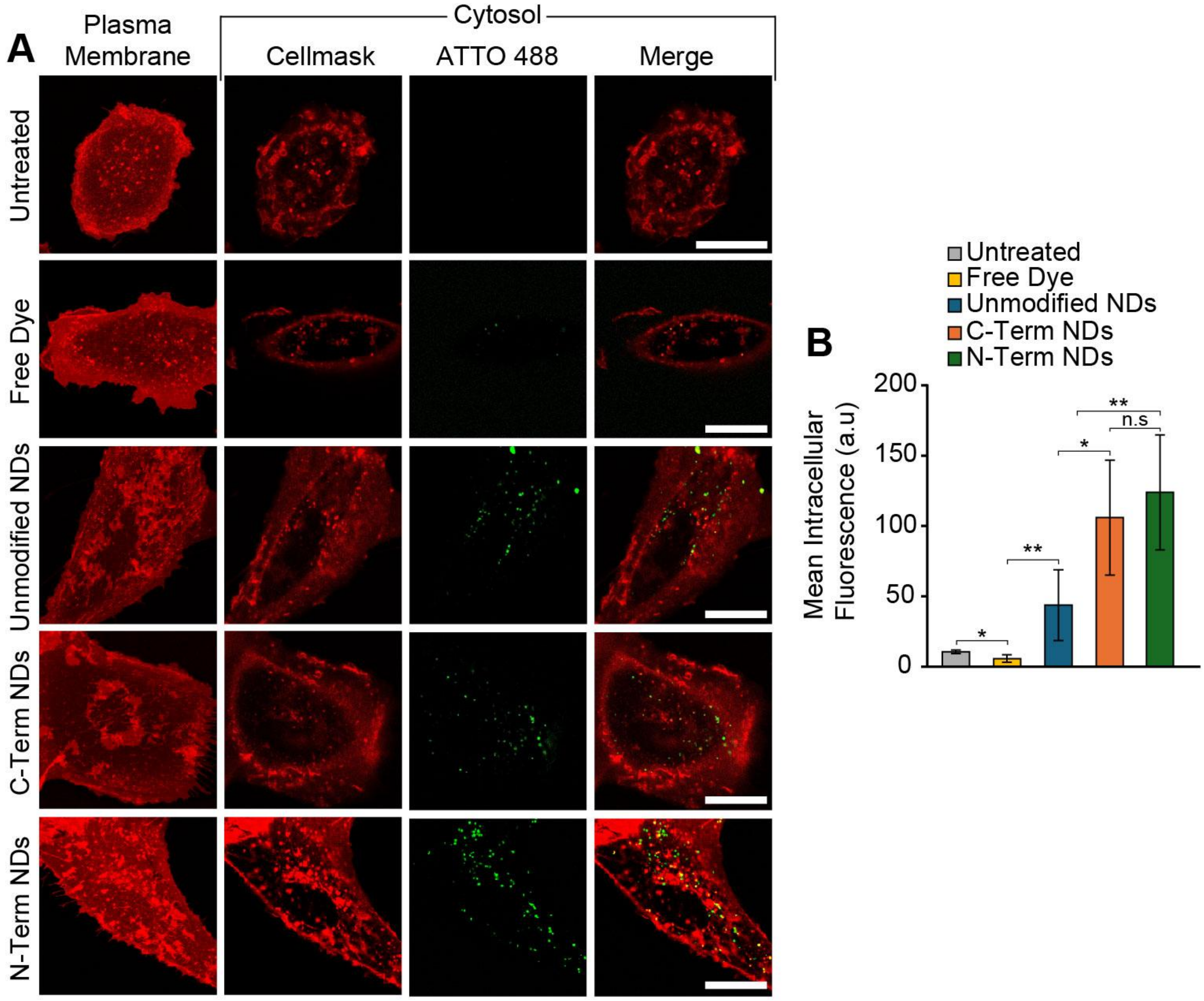


**Figure S9: R6W3-functionalized NDs undergo cellular internalization in RPE cells.** (A) Representative confocal images of RPE cells incubated with fluorescent NDs. (B) Quantification of intracellular fluorescence in RPE cells. Data are presented as mean ± SD. Statistical significance was determined by two tailed unpaired Student's t-test (* = p < .05, ** = p < .01, *** = p < .001, **** = p < .0001, n.s = not significant). Scale bars: 25 µm.

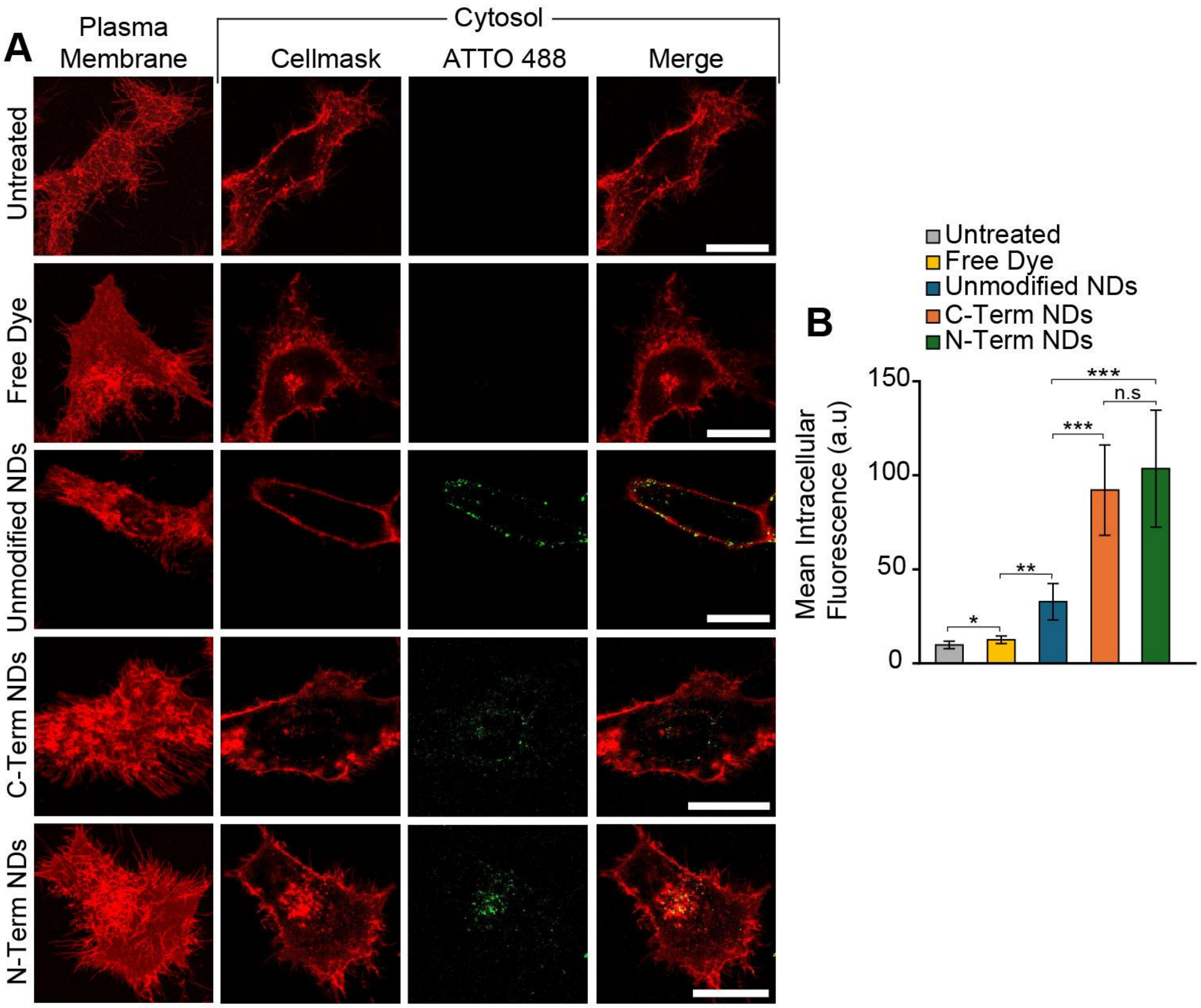


**Figure S10: R6W3-functionalized NDs undergo cellular internalization in HeLa cells.** (A) Representative confocal images of HeLa cells incubated with fluorescent NDs. (B) Quantification of intracellular fluorescence in HeLa cells. Data are presented as mean ± SD. Statistical significance was determined by two tailed unpaired Student's t-test (* = p < .05, ** = p < .01, *** = p < .001, **** = p < .0001, n.s = not significant). Scale bars: 25 μm.

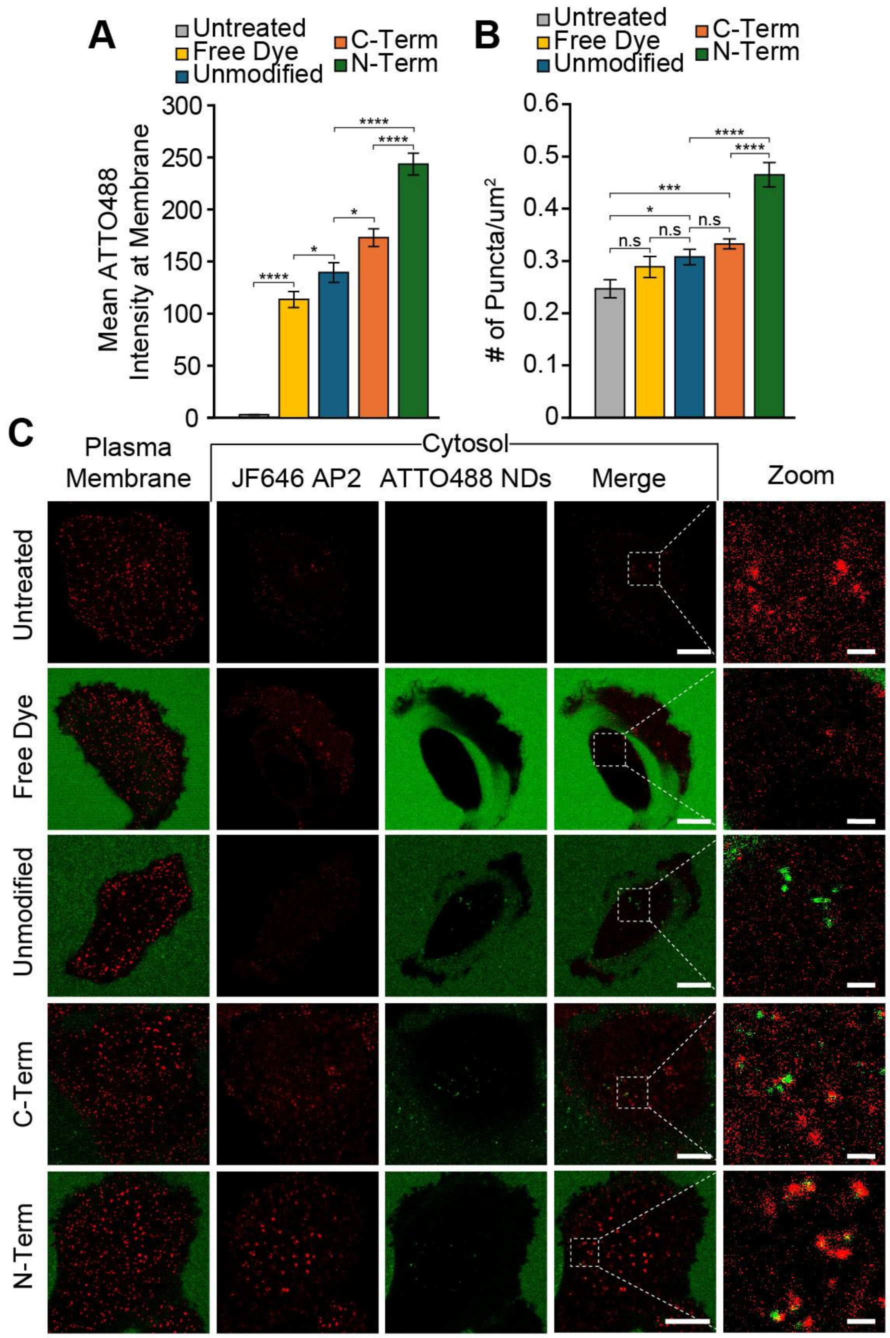


**Figure S11: R6W3-functionalized NDs enhance membrane association and puncta signal in SUM159 cells.** (A) Quantification of membrane-associated ATTO 488 signal in SUM159 cells. (B) Quantification of puncta density. (C) Confocal images of SUM159 cells expressing AP2-HaloTag labeled with JF646 (red) and incubated with ATTO488 labeled NDs (green). Insets show magnified regions of interest. Data are presented as mean ± SEM. Statistical significance was determined by two tailed unpaired Student's t-test (* = p < .05, ** = p < .01, *** = p < .001, **** = p < .0001). Scale bars: 10 μm (main images) and 1 μm (zoom).

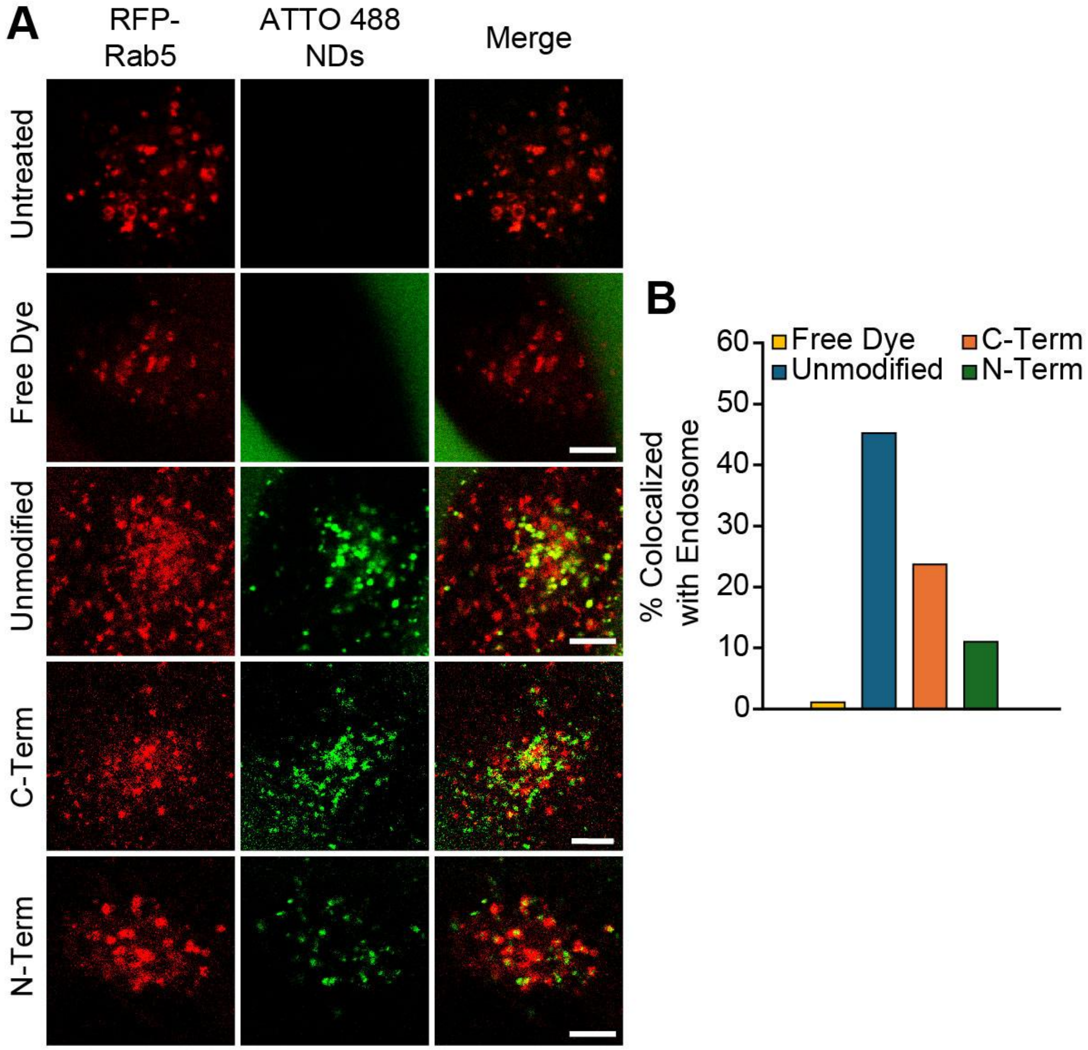


**Figure S12: R6W3-functionalized NDs exhibit reduced colocalization with early endosomes in SUM159 cells.** (A) Representative confocal images of SUM159 cells expressing Rab5-RFP (red) and incubated with ATTO 488 labeled NDs (green) or free dye. (B) Quantification of ATTO 488 colocalization with endosomes across treatment conditions. Scale bars = 5 μm.

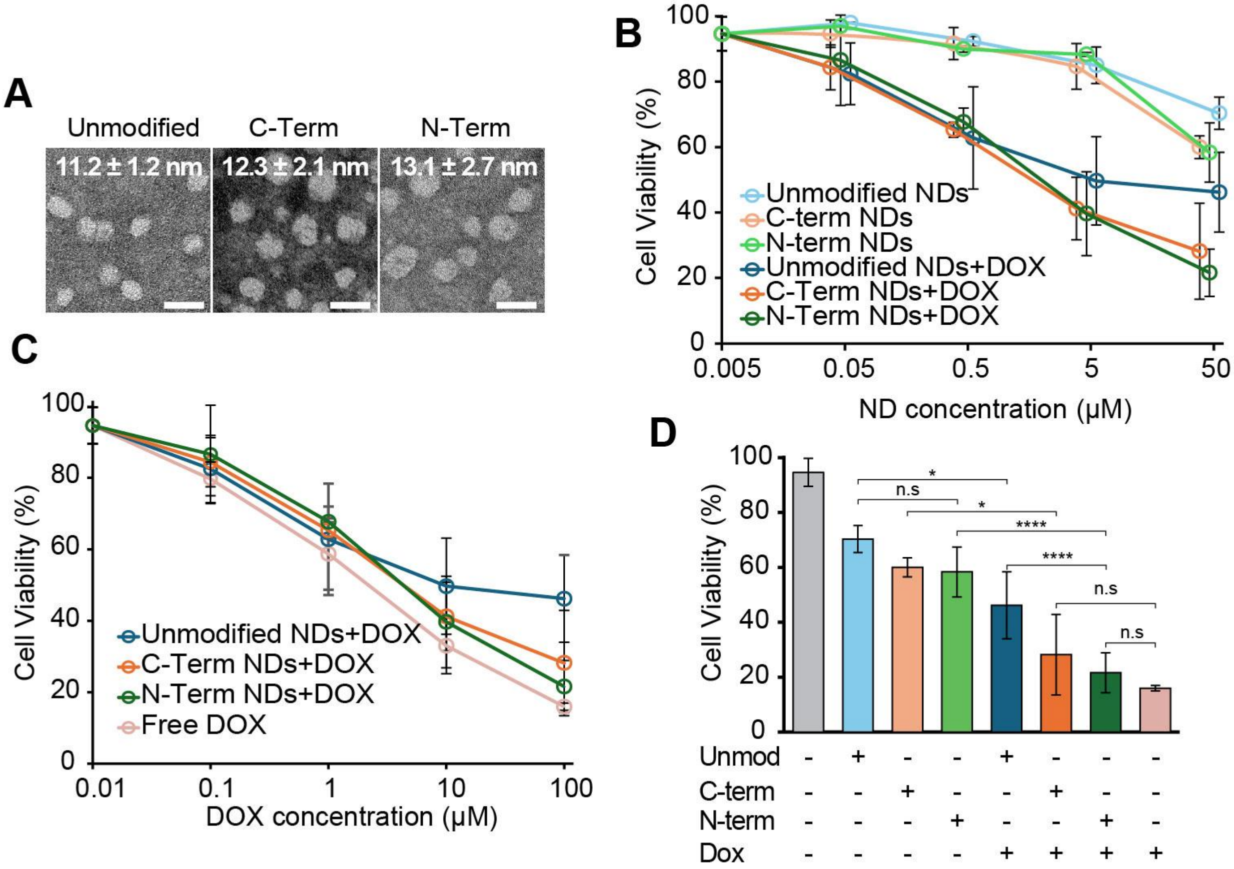


**Figure S13: DOX-loaded R6W3-functionalized NDs exhibit enhanced cytotoxicity in SUM159 cells.** (A) Representative TEM images of unmodified, C-term, and N-Term NDs. TEM size data are presented as mean ± SEM. (B) Cell viability as a function of ND concentration for unloaded and DOX-loaded NDs. (C) Cell viability as a function of DOX concentration. (D) Comparison of cell viability across treatment conditions at the highest DOX concentration tested. Cell viability data are presented as mean ± SD. Statistical significance was determined by two tailed unpaired Student's t-test (* = $p < .05$, ** = $p < .01$, *** = $p < .001$, **** = $p < .0001$, n.s = not significant). Scale bars: 20 nm.

Table S1: Protein Sequences

| | |
|---|---|
| MSP1D1 (Unmodified) | MGHHHHHHHDYDIPTTENLYFQGSTFSKLREQLGPVTQEFWDNLEK ETEGLRQEMSKDLEEVKAKVQPYLDDFQKKWQEEMELYRQKVEPL RAELQEGARQKLHELQEKLSPLGEEMRDRARAHVDALRTHLAPYSD ELRQRLAARLEALKENGGARLAEYHAKATEHLSTLSEKAKPALEDLR QGLLPVLESFKVSFLSALEEYTKKLNTQ |
| MSP1D1-R6W3 (C-Term) | MGHHHHHHHDYDIPTTENLYFQGSTFSKLREQLGPVTQEFWDNLEK ETEGLRQEMSKDLEEVKAKVQPYLDDFQKKWQEEMELYRQKVEPL RAELQEGARQKLHELQEKLSPLGEEMRDRARAHVDALRTHLAPYSD ELRQRLAARLEALKENGGARLAEYHAKATEHLSTLSEKAKPALEDLR QGLLPVLESFKVSFLSALEEYTKKLNTQGGGGSRRWWRRWRR |
| R6W3-MSP1D1 (N-term) | MRRWWRRWRRGGGGSDYDIPTTENLYFQGSTFSKLREQLGPVTQ EFWDNLEKETEGLRQEMSKDLEEVKAKVQPYLDDFQKKWQEEMEL YRQKVEPLRAELQEGARQKLHELQEKLSPLGEEMRDRARAHVDAL RTHLAPYSDELRQRLAARLEALKENGGARLAEYHAKATEHLSTLSEK AKPALEDLRQGLLPVLESFKVSFLSALEEYTKKLNTQGGGGSHHHH HH |
| eGFP (unmodified) | MRGSHHHHHHGMASMTGGQQMGRDLYDDDDKDRWGSMVSKGE ELFTGVVPILVELDGDVNGHKFSVSGEGEGDATYGKLTLKFICTTGKL PVPWPTLVTTLTYGVQCFSRYPDHMKQHDFFKSAMPEGYVQERTIF FKDDGNYKTRAEVKFEGDTLVNRIELKGIDFKEDGNILGHKLEYNYN SHNVYIMADKQKNGIKVNFKIRHNIEDGSVQLADHYQQNTPIGDGPV LLPDNHYLSTQSKLSKDPNEKRDHMVLLEFVTAAGITLGMDELYK |
| eGFP-R6W3 (C-Term) | MVSKGEELFTGVVPILVELDGDVNGHKFSVSGEGEGDATYGKLTLKF ICTTGKLPVPWPTLVTTLTYGVQCFSRYPDHMKQHDFFKSAMPEGY VQERTIFFKDDGNYKTRAEVKFEGDTLVNRIELKGIDFKEDGNILGHK LEYNYNSHNVYIMADKQKNGIKVNFKIRHNIEDGSVQLADHYQQNTP IGDGPVLLPDNHYLSTQSKLSKDPNEKRDHMVLLEFVTAAGITLGMD ELYKGGGGSRRWWRRWRR |
| R6W3-eGFP (N-Term) | MRRWWRRWRRGGGGSGMASMTGGQQMGRDLYDDDDKDRWGS MVSKGEELFTGVVPILVELDGDVNGHKFSVSGEGEGDATYGKLTLKF ICTTGKLPVPWPTLVTTLTYGVQCFSRYPDHMKQHDFFKSAMPEGY VQERTIFFKDDGNYKTRAEVKFEGDTLVNRIELKGIDFKEDGNILGHK LEYNYNSHNVYIMADKQKNGIKVNFKIRHNIEDGSVQLADHYQQNTP IGDGPVLLPDNHYLSTQSKLSKDPNEKRDHMVLLEFVTAAGITLGMD ELYKGGGGSHHHHHH |
